# Four numbers, one axis: deep learning models reveal what leaf spectrum constrains about Farquhar-von Caemmerer-Berry photosynthesis

**DOI:** 10.64898/2026.08.27.747677

**Authors:** Rishav Ray, Julin N. Maloof, Troy S. Magney

## Abstract

- Leaf reflectance spectra are emerging as a viable substitute for gas-exchange measurements of photosynthetic capacity, with a community benchmark reporting that a spectrum accurately recovers most Farquhar–von Caemmerer–Berry (FvCB) parameters.
- This study re-scores the recovery under dataset-blocked, species-blocked, and leave-one-dataset-out designs, measuring the split-half reliability of each curated parameter. We constructed a convolutional encoder that maps a spectrum to the four parameters through a fixed, differentiable FvCB decoder trained on measured assimilation.
- A conspecific of 97.4% of held-out leaves were present in the training set, and accuracy is lost along the dataset axis but not along the species axis. Under blocked evaluation, a spectrum constrains a single capacity axis. Jmax25 retains only 17% of its recovery when Vcmax25 is held constant, and the Jmax25:Vcmax25 ratio is not predicted above a median null. The curated values of TPU25 are not reproducible, whereas those of Rday25 are well determined, but its recovery fails due to the loss.
- The published study measures interpolation rather than transfer, and spectra constrain less of the FvCB parameter space than assumed, including the carboxylation to electron transport balance. Routing predictions through explicit biochemistry makes identifiability measurable, although it does not improve prediction accuracy.

## Introduction

Terrestrial photosynthesis removes approximately 120 petagrams of carbon from the atmosphere annually, which is an order of magnitude greater than annual fossil-fuel emissions. The persistence of this uptake under a warming climate depends on the accuracy with which Earth system models represent the biochemistry of individual leaves (Beer *et al*., 2010). These models utilize the Farquhar–von Caemmerer–Berry (FvCB) model, where net assimilation is the minimum of three potential rates, that are, rubisco carboxylation, electron transport, and triose phosphate utilization (Farquhar *et al*., 1980). Four temperature-normalized parameters determine these rates for a given leaf, maximum carboxylation capacity (Vcmax25), maximum electron transport capacity (Jmax25), maximum rate of triose phosphate utilization (TPU25), and day respiration (Rday25). Uncertainty in Vcmax25 is a leading source of error in modelled carbon uptake at regional and global scales (Rogers, 2014; Rogers *et al*., 2017; Busch *et al*., 2024). Additionally, the Jmax25:Vcmax25 ratio must vary with climate and acclimation state rather than holding at a fixed value (Kattge & Knorr, 2007; Smith *et al*., 2019). The conventional method for determining all four parameters is the A-C_i_ curve, which involves measuring net assimilation across a range of intercellular CO2 concentrations and fitting the parameters simultaneously. However, a single curve requires 30 to 60 minutes of measurement time, making large sampling across species, sites, and growing seasons impractical (Stinziano *et al*., 2019). Leaf reflectance spectra offer a faster alternative, as they can be collected in seconds. Vegetation traits are then retrieved from spectra using empirical regression, inversion of physically based radiative transfer models, or hybrid schemes that train a statistical model on radiative transfer simulations (Jacquemoud & Baret, 1990; Verrelst *et al*., 2019). For photosynthetic capacity, the empirical route has been dominant where partial least squares regression (PLSR) models trained on paired spectra and gas-exchange measurements have been used to predict Vcmax and Jmax since Serbin *et al*. (Serbin *et al*., 2012). This approach has been extended across crops and wild species with reported accuracies of R^2^ ≈ 0.5–0.8 (Yendrek *et al*., 2017; Fu *et al*., 2019; Meacham-Hensold *et al*., 2019; Cui *et al*., 2025), and are supported by an established set of community best practices (Burnett *et al*., 2021). The Global Spectra-Trait Initiative (GSTI) compiles this work into a database of paired leaf spectra and curated FvCB parameters spanning multiple contributing datasets, along with a published benchmark reporting that a spectrum recovers most FvCB parameters with practically useful accuracy (Lamour *et al*., 2026).

This type of benchmark number is dependent on two factors, the design that produced it and the target against which it was scored. When observations are randomly split, any structural link among them, such as shared species, site, instrument, or growing conditions allows information from a held-out observation to reach the model through another observation of the same unit. This results in the benchmark describing recognition of a previously seen unit rather than prediction of an unseen one (Roberts *et al*., 2017; Kapoor & Narayanan, 2023; Stock *et al*., 2023). Random splits inflate reported accuracy compared to grouped or blocked splits in species distribution modeling (Roberts *et al*., 2017; Valavi *et al*., 2019), large-scale ecological mapping (Ploton *et al*., 2020), and in cases where instrument, operator and protocol act as grouping variables in their own right (Leek *et al*., 2010). The same dependence holds in spectroscopy, where the validation method (random or shift-aware split) determines which approach appears superior (Passos, 2026). Domain shift between campaigns, sites, and sensors is now recognized as the central obstacle to deploying spectral models beyond their training data (Ma *et al*., 2024), and has been characterized directly for foliar trait retrieval (Huang *et al*., 2026). In a recent multi-season study of PLSR-based trait prediction in maize, genotype-exclusive folds were required to prevent leakage between calibration and validation sets (Xu *et al*., 2026). GSTI’s published design involves a random split stratified within each contributing dataset. Therefore, every held-out leaf is drawn from a dataset also present in the training set. Since many contributing datasets sample the same species at multiple growth stages, treatments, or timepoints, a held-out leaf’s own species is frequently present in the training set as well. On the other hand, the target is the curated FvCB parameters, which are fitted estimates rather than direct measurements. Each parameter is identifiable only where a curve samples the C_i_ region that constrains it. TPU25, in particular, is identifiable only from curves that reach the TPU-limited region at high C*_i_* (Gu *et al*., 2010; Sharkey, 2019). Fitted values depend on the choice of fitting tool and its assumptions (Lochocki *et al*., 2025; Wei *et al*., 2026). A correlation against a target of limited reproducibility is bounded by that reproducibility, irrespective of the model that produced it.

To determine what a spectrum constrains parameter by parameter rather than in aggregate, it is necessary to use a model whose predictions are directly influenced by the parameters themselves. Deep learning has become standard practice in remote sensing (Zhu *et al*., 2017; Ma *et al*., 2024), including for hyperspectral data, where convolutional and attention-based architectures learn directly from contiguous bands rather than selected indices (Audebert *et al*., 2019; Paoletti *et al*., 2019; Hong *et al*., 2022). Additionally, large self-supervised models pretrained on spectral imagery are being transferred to downstream tasks as well (Hong *et al*., 2024). One-dimensional convolutional networks applied to continuous spectra match or exceed PLSR in chemometric calibration and learn an appropriate spectral filter instead of relying on hand-specified preprocessing steps (Cui & Fearn, 2018; Ng *et al*., 2019). Both convolutional and attention-based models have been applied to leaf traits and photosynthetic capacity, with mixed results compared to PLSR (Song & Wang, 2021; Furbank *et al*., 2021; Deng *et al*., 2025). These models are trained in the same manner as PLSR, directly against each fitted parameter and scored on how closely they reproduce its value. This approach does not distinguish between a spectrum that constrains the parameter and a model exploiting incidental covariance among traits in the training data (Kothari & Schweiger, 2022). Routing predictions through an explicit physiological model addresses this issue by allowing each parameter to be perturbed while others are held fixed, and the effect on predicted assimilation is directly read. This process mirrors standard operations in practical identifiability analysis (Raue *et al*., 2009). Differentiable hybrid modeling provides this exact machinery. Embedding a neural network within a process model and training the two jointly by propagating gradients through both has been used to learn unconstrained parameters and process representations in hydrology and land-surface modeling while maintaining outputs tied to known physical structure (Reichstein *et al*., 2019; Rackauckas *et al*., 2021; Shen *et al*., 2023). Photosynthesis is an established case within this lineage. A fully differentiable implementation of the FATES photosynthesis routine learns Vcmax25 and related parameters from environmental covariates rather than fitting them site by site (Aboelyazeed *et al*., 2023), and has since been extended to parameter acclimation (Aboelyazeed *et al*., 2025). Related frameworks couple neural networks to physical representations of stomatal and aerodynamic resistance (ElGhawi *et al*., 2023) and of soil water limitation on carbon and water fluxes (Fang & Gentine, 2024). These architectures are frequently credited with improving prediction. To validate this claim, a control is needed where the physical constraint is removed and everything else held fixed. Often, reported advantages of physics-informed models do not withstand stronger controls (McGreivy & Hakim, 2024).

In this study, we integrate a one-dimensional convolutional encoder with a fixed, differentiable FvCB decoder to ensure that spectral predictions pass through the same four parameters that the gas-exchange community fits and reports. We train this architecture against measured assimilation rather than pre-fitted parameter values. This approach allows us to address four sequential questions. First, what shared structure, such as dataset or species, exists across the train/test boundary in the published benchmark split, and how does it impact accuracy under a stricter design. Second, what specific information does the gas-exchange curve identify, and how reproducible are the curated parameters against which any spectral model is scored. Third, given the identified references, which parameters does a reflectance spectrum constrain once leakage is removed, and are these constraints independent of one another. Fourth, What is the cost in prediction accuracy when routing spectral predictions through fixed biochemistry, and what benefits does it provide? Is the constraint itself responsible for any observed advantages in the architecture’s performance.

## Methods

### Dataset

We utilized the Global Spectra-Trait Initiative (GSTI) database (https://github.com/plantphys/gsti), which pairs leaf reflectance spectra with A-C_i_ gas-exchange curves and curated Farquhar–von Caemmerer–Berry (FvCB) parameters across contributed datasets. We restricted our analysis to C3 species and removed spectra with negative reflectance values, leaving 7549 of the original 7561 curated observations. From these, we defined two cohorts. Cohort P (paired: 2983 curves, 23 datasets, 230 species) comprises curves with a full-range spectrum, a complete A-C_i_ curve, and GSTI-fitted parameters. This cohort serves as the basis for all primary tier comparisons. Cohort E (extended: 4911 curves) includes one-point measurements and curves with partial or discarded GSTI fits, and is used only where explicitly noted as the Cohort E training control. Among the 2983 curves in Cohort P, 42.6% correspond to crop species. Cohort P’s 2983 curves differ from the roughly 2443 curves and 36 datasets reported in the original GSTI publication. This discrepancy is due to the additions made in 2026, after the original publication. We retained these new entries rather than reconstructing the earlier database state.

### Decoder and decoder validation

We implemented the FvCB model as a fixed, non-trainable decoder that maps four leaf parameters (Vcmax25, Jmax25, TPU25, Rday25), along with measured C_i_, Tleaf, and Qin, to the predicted net assimilation A. Gross assimilation is determined as the co-limited minimum of three potential rates – Rubisco-limited (Ac), electron-transport-limited (Aj), and triose-phosphate-utilization-limited (Ap) – combined pairwise through a non-rectangular hyperbola with θ_cj_ = θ_ip_ = 0.999. Electron transport J is computed from Jmax using a non-rectangular hyperbola with absorptance 0.85, apparent quantum yield 0.425, and curvature Θ = 0.7. TPU-limited capacity is fixed at Wp = 3 × TPU rather than fitted independently, and we used the larger root of the co-limitation quadratic whenever C_i_ ≤ the CO2 compensation point Γ, following GSTI’s own implementation. Day respiration Rday is subtracted once from the co-limited gross rate. Temperature scaling employs GSTI’s Arrhenius and peaked-Arrhenius constants (Bernacchi, Harley, and Sharkey parameterizations as implemented in GSTI’s code), with a +273 °C-to-K conversion and reference temperature TRef = 298.16 K. We reproduced these to ensure that decoder output remains directly comparable to GSTI’s curated values.

We validated the decoder by fitting all four FvCB parameters directly to each curve’s own measured points (Tier 0; Fig. 1, see below), independently within each of GSTI’s four curve model classes (Ac, Ac_Aj, Ac_Aj_Ap, Ac_Ap), and comparing the fitted values to the curated values (Table S2). Across all 2983 Cohort P curves, the fitted and curated values agreed with a slope of 1.004 and an R^2^ of 0.998. Within each model class, the median fit RMSE matched GSTI’s own per-curve residual σ almost exactly (RMSE/σ = 0.9996–1.0000). The decoder’s free fit converges to essentially the same optimum that GSTI’s own fitter finds for each curve, which represents the ceiling of agreement rather than a shortfall.

**Fig. 1.**
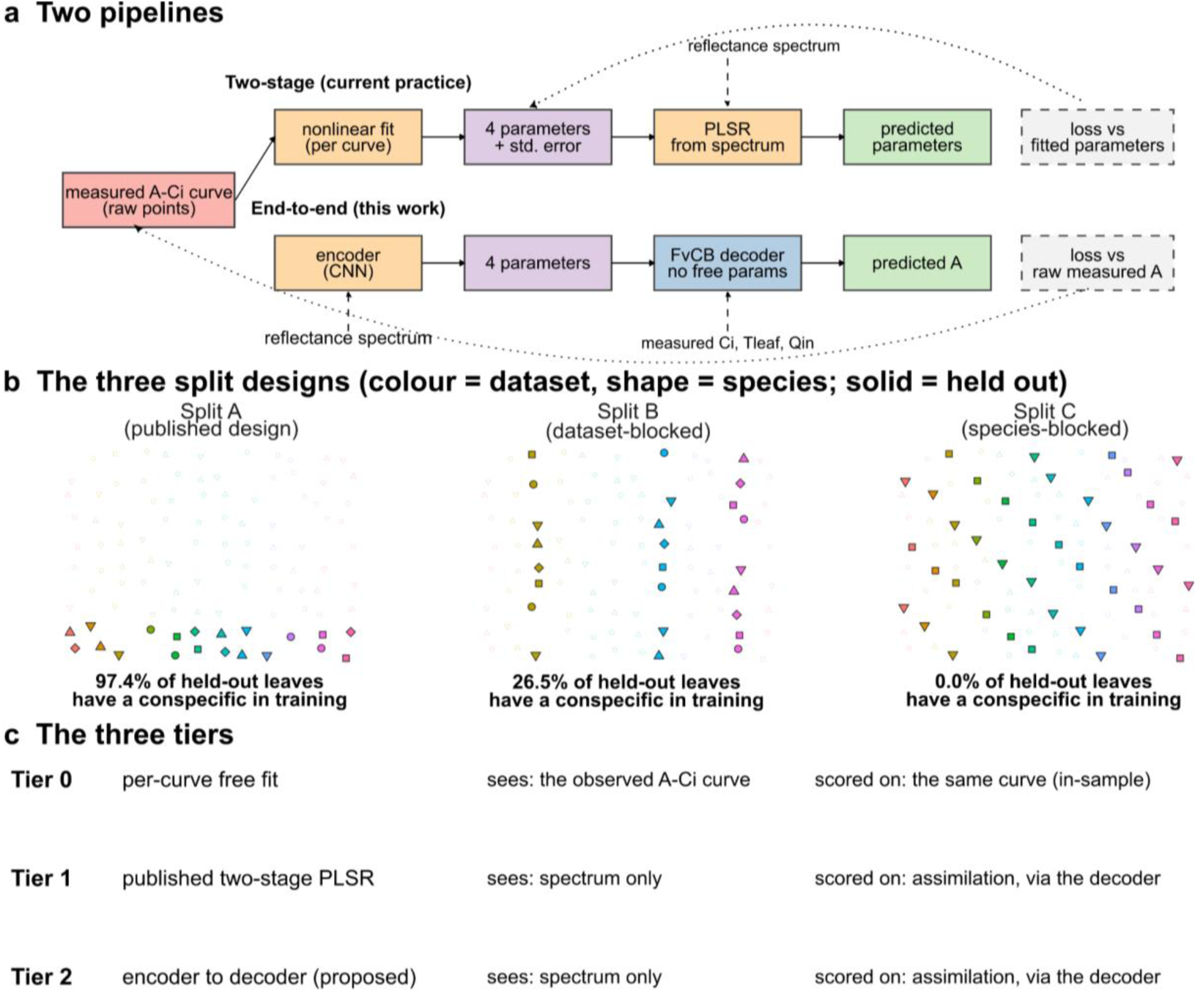
Study design. a) The two pipelines and what each is scored against. The two-stage baseline (Tier 1) fits FvCB parameters to each gas-exchange curve and then regresses spectra on those fitted parameters; the constrained model (Tier 2) predicts the four parameters directly from the spectrum and pushes them through a differentiable FvCB decoder, so its loss is computed in assimilation space against the measured curve. b) The three split designs (schematic; colour = dataset, shape = species). Split A is the published design, a random 80/20 split stratified within dataset. Split B assigns whole datasets to folds; Split C assigns whole species to folds. c) The three tiers, Tier 0 (curve-level free fit, in-sample), Tier 1 (two-stage baseline), Tier 2 (spectrum → parameters → decoder).

### Tiers

We compared three tiers that differ in what they observe and what they are scored against. Tier 0 is a per-curve free fit, where the four FvCB parameters are optimized to match a single curve’s own measured points, with no spectrum involved, no parameters shared across curves, and no held-out evaluation. This tier serves to validate the decoder (as described above) and, later, to measure how much information a curve alone carries about each parameter (the thinning experiment, below). Since it is fit and scored in-sample on the same curve, Tier 0 acts as a decoder-adequacy reference rather than a ceiling on what a spectrum-only model could achieve. Tier 1 is GSTI’s published two-stage pipeline, transcribed and unchanged where FvCB parameters are fitted per curve and then predicted from the spectrum using partial least squares regression (PLSR). Tier 2 is the architecture under study in which an encoder maps the spectrum directly to the four FvCB parameters, which are then passed through the fixed decoder described above. The loss is computed against raw measured assimilation rather than against fitted parameters.

### Encoder and training

The Tier 2 encoder is a four-block one-dimensional convolutional network (channel widths 32-64-128-128, kernel size 7) over two input channels, reflectance and its first derivative, each vector-normalized per observation. Each block is followed by batch normalization, ReLU activation, and max pooling. The network architecture includes global average pooling and a shared 64-unit multi-layer perceptron (MLP) layer. Two linear heads read from this shared representation, a four-unit head producing Vcmax25, Jmax25, TPU25, and Rday25, which are passed to the decoder, and a separate one-unit auxiliary head producing Rdark. The auxiliary head is trained against measured dark respiration but does not enter the decoder. Rday and Rdark represent distinct physiological quantities, such as respiration in light, inferred from the gas-exchange curve, versus respiration measured in darkness and collapsing them into a single head would conflate two different measurements.

Both heads employ a softplus activation, ensuring predictions remain positive. This aligns with the physical positivity of FvCB parameters and Rday’s near-positivity. Importantly, this is a genuine constraint where approximately 9% of curated Rday25 values in Cohort P are negative, making them unattainable by construction. Each head’s output is further scaled multiplicatively against the training-set median value for that parameter (prediction = median × softplus (head output + c), with c fixed so that softplus(c) = 1 at initialization). This ensures training begins at the median parameter values rather than an arbitrary point on the decoder’s response surface.

The training loss is a σ-weighted Gaussian negative log-likelihood on assimilation, using GSTI’s curated per-curve residual σ as the weight. This loss is summed over valid points within a curve and then averaged across curves in a batch. Summing within a curve before averaging across curves ensures that each curve’s total contribution is proportional to the information it carries, automatically down-weighting one-point curves relative to full curves without requiring a separate estimator for measurement count. A fixed auxiliary MSE term on the Rdark head is added with weight λaux = 0.1, set in advance and never tuned. Training was conducted using Adam optimizer (learning rate 1 × 10^-3^, batch size 64) for up to 300 epochs with early stopping (patience set at 20 epochs on validation loss). Each configuration was trained across 10 random seeds, and ensemble statistics (mean and seed-to-seed standard deviation) are reported throughout.

### Splits

We used five evaluation designs. Split A is GSTI’s published design that is a random 80/20 split stratified within each dataset, so held-out leaves are drawn from datasets also present in training. Splits B and C are grouped splits where Split B assigns entire datasets to 5 folds (684, 572, 575, 576, 576 curves), and Split C assigns entire species to 5 folds (596–597 curves each, 41–48 species per fold), so that no held-out dataset or species appears in that fold’s training portion. Splits BE and CE are Cohort E variants of B and C, adding Cohort E’s extra curves to the training portion of each fold while keeping the same held-out curves. Leave-one-dataset-out (LODO) holds out each of the 17 Cohort P datasets with at least 40 curves individually (ten seeds each), with the remaining six smaller datasets pooled into one additional held-out block reported separately.

Split A’s 589-curve test partition was frozen when splits were constructed and was not read, inspected, or used in any decision until the designated evaluation checkpoint, after all tiers, architectures, and hyperparameters were finalized. Because that partition is drawn from Split A, however, it inherits the within-dataset design’s leakage, and we therefore report parameter recovery under Splits B and C throughout, carrying the frozen-set values only as an explicitly labelled leaky reference. Freezing protects against adaptive overfitting to the test set, but it does not correct a split design.

Split A therefore yields two distinct scores, and we report both. The first is computed on the frozen 589-curve test partition. The second is computed by inner five-fold cross-validation over Split A’s 2394 remaining curves, where those curves are divided at random into five folds, each fold is predicted by a model trained on the other four, and the frozen partition is excluded from both training and prediction throughout. Both tiers use this identical construction, so the two are directly comparable. The inner-cross-validation score exists because Splits B and C are themselves scored by cross-validation over all 2983 curves, and comparing the three designs requires a Split A score computed the same way rather than one restricted to a 589-curve subset.

### Metrics

We report skill rather than R^2^ as the primary metric. Skill = 1 − (RMSEmodel / RMSEnull), where the null prediction, for a given held-out subset, decodes each held-out curve’s points using the median FvCB parameters of that subset’s training portion, ignoring the spectrum entirely. Skill = 0 is exact parity with this subset-specific null; skill < 0 means the model performs worse than the training-portion median. Because the null is recomputed separately for every subset, each LODO fold, each split, each stratum, skill values are comparable across subsets of different size and composition in a way R^2^ is not, since R^2^’s denominator is the variance of whichever curves happen to fall in that particular held-out set. We report skill in assimilation space (predicted versus measured A) throughout, and note explicitly wherever a reported R^2^ is instead in parameter space (predicted versus curated FvCB parameters)..

### Thinning experiment

To measure how much information an A-C_i_ curve carries about each FvCB parameter, independent of any spectral model, we thinned each curve to a fixed coverage (8, 6, 4, 2, or 1 of the original points retained) under three removal schemes. Uniform (points dropped without regard to C_i_), high-C_i_-dropped (highest-C_i_ points removed first), and low-C_i_-dropped (lowest-C_i_ points removed first). We refit Tier 0 to each thinned curve and compare the resulting parameters to the same curve’s full-coverage fit. Coverage of 4 or fewer points is flagged as underdetermined and excluded from the degradation comparisons, because a four-parameter model can exactly interpolate 4 or fewer points regardless of where they fall, making the fit an artifact of the optimizer’s starting values rather than a measurement of the curve.

### Target reliability

Correlations against a curated target are bounded by how reproducible that target is, so we measured the reproducibility of GSTI’s own fitted parameters before interpreting any prediction failure. For each curve with at least 6 measured points, we split the points into odd- and even-indexed halves, fitted all four FvCB parameters independently to each half by the same Tier 0 free fit used for decoder validation, and correlated the two resulting estimates across curves. Because each half carries only half the points, this understates the reliability of a full-curve fit, so we applied the Spearman–Brown correction r_SB = 2r/(1 + r) and reported the ceiling on any predictor as √r_SB. Both the raw and corrected values are reported, and the correction is applied only where r > 0. Curves whose fit failed to converge in either half were excluded from that parameter’s correlation. We report reliability database-wide, within each GSTI model class, and as a function of curve coverage, both for the subset of curves with at least 12 points, so that each half carries at least six, and across coverage bins (8, 9–11, 12–15, and 16 or more points).

Two diagnostics accompany the Pearson correlation. We report Spearman rank correlation alongside it, because a fitter bound produces a cluster of identical extreme values that acts as high leverage on a Pearson coefficient, and we report the fraction of curves with at least one half-fit pinned at a parameter bound. Where those two diagnostics disagree with the Pearson value we report the bound-excluded correlation as well, and treat the bound-hitting rate rather than the correlation as the primary statement about that parameter. We applied the same odd/even procedure to the Jmax25:Vcmax25 ratio, computing the ratio within each half-fit and correlating across curves, to establish that the ratio is a reproducible target before testing whether it can be predicted.

### Parameter independence

Two parameters can each correlate with their curated values while carrying only one underlying axis of variation. We tested this three ways, none of which is sufficient alone. First, partial correlation, where we regressed both predicted and curated Jmax25 (and TPU25) on curated Vcmax25 by ordinary least squares, correlated the residuals, and reported the fraction of the raw correlation retained. Second, ratio prediction, where we scored the predicted Jmax25:Vcmax25 ratio against a null equal to the median curated ratio of the training portion, the ratio analogue of the median-parameter null used throughout. Because unbounded linear extrapolation can drive a predicted denominator arbitrarily close to zero, ratios were computed only where predicted Vcmax25 was at or above the fitter’s own lower bound of 1.0 µmol m^-2^ s^-1^. Third, and needing no curated value at prediction time, we correlated each method’s own predicted Vcmax25 against its own predicted Jmax25 and compared that to the correlation between the curated values themselves. All three were run for both tiers, so that a collapse specific to the constrained architecture can be distinguished from one belonging to leaf spectra generally.

We also quantified range compression by regressing predicted on curated values and reporting the slope, and by binning held-out curves into curated deciles and reporting the mean signed bias per decile as a percentage of that decile’s own curated mean, which puts parameters with different units on a common scale.

### Loss-surface diagnostics

To ask whether a prediction failure is architectural rather than biological, we probed the decoder’s loss surface directly. For each parameter in turn we perturbed the predicted value on held-out curves and recomputed the assimilation negative log-likelihood, reporting the change as a fraction of the unperturbed value. The perturbation is 10% of that parameter’s own interquartile range across Cohort P rather than a fixed absolute step, because the four parameters span two orders of magnitude and a common absolute step would not pose them the same question. An absolute ±2 µmol m^-2^ s^-1^ step is additionally reported for Rday25 and is not comparable to the others.

We separately computed the analytic sensitivity of decoded assimilation to each parameter by automatic differentiation through the decoder, evaluated point by point along the C*i* axis rather than on a summed loss, since summing collapses the axis of interest. This was done both on a dense C*i* grid at Cohort P’s median parameter vector and at every measured point of each fully parameterised curve using that curve’s own parameters, temperature, and irradiance. We report the within-curve variability of each derivative, which distinguishes a parameter whose influence is concentrated in a specific C_i_ region from one whose influence is uniform along the curve and therefore carries no positional information for the fitter to exploit.

Where a parameter reached through the decoder failed, we compared it against a directly supervised head on the same encoder and the same splits, which isolates the routing of the target from the information content of the spectrum. Two further checks were run on such failures. The ratio of predicted to curated variance on held-out curves, to test whether the head had collapsed to a near-constant, and the correlation between that parameter’s residuals and the residuals of the other three, to test whether error was being absorbed across parameters.

### Alternative-explanation controls

The transfer comparison between tiers admits several explanations besides the physics constraint, and we tested each against a separate control. Each control manipulates one factor across a stated range and is judged against the skill gap it would need to explain.

Tier 1 fits a separate PLSR per trait and can use only curves for which that trait was individually fitted, whereas Tier 2 trains jointly against raw assimilation and uses every curve carrying at least one parameter. We therefore retrained Tier 2 from scratch on only the curves Tier 1 could use for each trait, one restricted model per trait, replicating Tier 1’s own per-trait inclusion rule.

Only the training set was restricted whereas the held-out population was never restricted. Therefore, all restricted models are scored on identical curves and the comparison is not confounded by a change in what is being predicted. Each restricted model is scored on its own trait under leave-one-dataset-out at the same seed count as the unrestricted model.

We fitted Tier 1’s own PLSR algorithm, unchanged including its component-selection procedure, on Tier 2’s input representation (5 nm binning, per-observation vector normalization, added first-derivative channel) in place of GSTI’s, and scored it in parameter space under both blocked designs. Component counts were re-selected under each preprocessing rather than carried over, since the optimal number depends on the input representation. The mean change is taken over the four FvCB parameters and Rdark25 is computed and reported but excluded from that mean.

We trained an otherwise identical encoder with the decoder removed, regressing directly on curated parameters under MSE, and evaluated it under leave-one-dataset-out at the same ten seeds as Tier 2. The design under which Tier 2’s advantage is largest and therefore the one with the most to explain. This control is what the sensitivity grid cannot provide, since every grid arm retains the decoder.

### Statistical comparisons

Paired comparisons between tiers under leave-one-dataset-out are Wilcoxon signed-rank tests at the dataset level, with each dataset contributing one ensemble mean per method (n = 17). We pair at the dataset level rather than the seed level because Tier 1 is a single deterministic fit per fold with no seed ensemble. Pairing across seeds would reuse each of its 17 values ten times and inflate the apparent sample size to 170 without adding information. The pooled block of six datasets with fewer than 40 curves is plotted for completeness but excluded from every paired test. As a distribution-free check we ran a cluster permutation test that flips the sign of each dataset’s paired difference across 20,000 permutations, which uses the per-seed information without treating seeds as independent replicates; it agrees with the dataset-level signed-rank test throughout.

Because several contributing datasets share a first author and probably an instrument and protocol, we repeated the paired test with same-lab datasets collapsed into 14 evaluation units. This removes same-lab duplication from the evaluation units but not from the training sets, since each dataset’s run retained its same-lab siblings in training.

To ask whether two models are equivalent rather than whether they differ we used two one-sided tests against a margin of ±0.033 fixed in advance at half the gap separating either network from Tier 1. We report the outcome of that test as specified even where it does not clear the margin, rather than substituting a post-hoc margin that would.

### Sensitivity grid

After the primary Tier 1/Tier 2 comparison under Split B was obtained, we ran a sensitivity grid to test whether the result depended on a specific, untuned choice of architecture. The grid varied encoder depth, channel width, kernel size, spectral binning, and the auxiliary loss weight, and was run only after the primary configuration was locked. It was not used to select the reported configuration. One grid arm, 1 nm spectral binning changes the input representation itself rather than encoder capacity, and we grouped it with a separate GSTI-preprocessing control rather than the architecture-only arms. Its Split B skill (0.214) falls below the Tier 1 baseline (0.282), which would otherwise misrepresent the range of outcomes achievable by architecture choice alone. Full arm-by-arm values, including the excluded binning arm and the control, are reported in Table S1.

### Implementation, code and data availability

The differentiable decoder, the convolutional encoder and all training and evaluation code were implemented in Python 3.11 using PyTorch 2.2.2, with NumPy and SciPy for the per-curve free fits and the statistical comparisons. The decoder is implemented in the same autograd framework as the encoder, which is what allows gradients from the assimilation loss to propagate through the fixed biochemistry into the encoder weights. The two-stage baseline is GSTI’s published PLSR pipeline, transcribed from its original R implementation. Figures were produced in R 4.2.1 using ggplot2. The source code and documentation is available at https://github.com/rishavray/dl_spectra_photosynthesis.

## Results

### The published design measures interpolation, not transfer

We first asked whether GSTI’s reported accuracy describes transfer to new unseen data, and if accuracy falls under a stricter split, which kind of shared structure is responsible. To that end, we counted what the published split shares across the train/test boundary and re-scored the same pipeline under two blocked designs and set the two blocking schemes against each other. Under the published split (Split A), 97.4% of held-out leaves have at least one conspecific already in the training set, whereas dataset-blocked holdout (Split B) reduces this to 26.5% and species-blocked holdout (Split C) to 0% by construction (Fig. 2a). All five traits recover close to their originally reported values under Split A, and R^2^ falls for every trait under Split B, with both respiration terms crossing zero (Rday25 from 0.60 to −0.33; Rdark25 from 0.59 to −0.23; Fig. 2b). Conspecific presence is not what drives that loss. Split C removes conspecific overlap entirely, yet recovers four of the five traits better than Split B. Partitioning held-out curves within the dataset-blocked folds by whether their own species appears in training changes skill by 0.006 (Fig. 2b). Holding out individually each of the 17 contributing datasets with at least 40 curves returns negative skill for four of them (Fig. 2c). Taken together, the published benchmark values describe interpolation within a dataset already represented in training, and the accuracy lost under a stricter split is lost along the dataset axis.

**Fig. 2.**
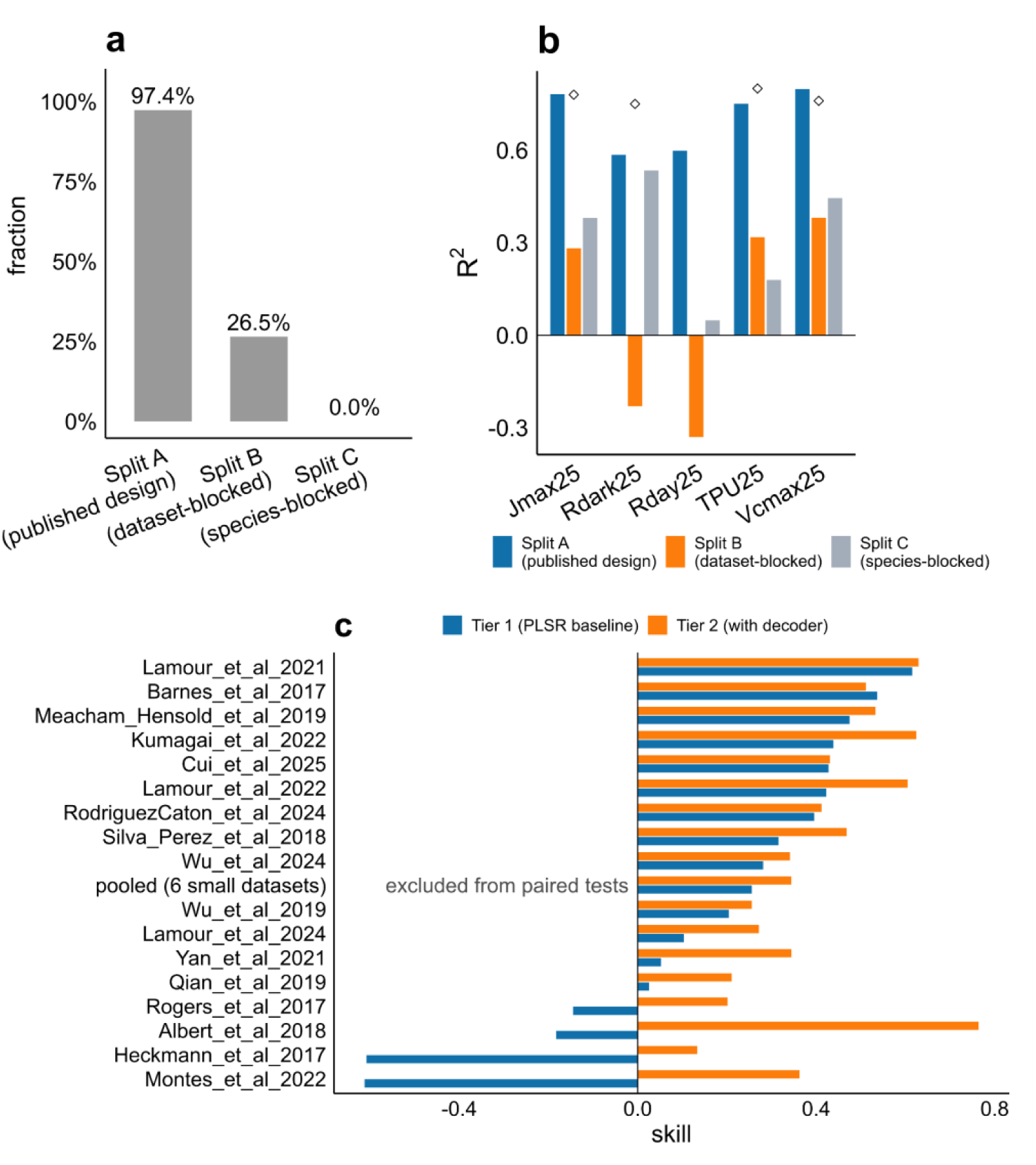
Benchmark of the published design under various splits. a) Fraction of held-out leaves that have a conspecific curve in the training partition, 97.4% under Split A, 26.5% under Split B, 0.0% under Split C. b) Parameter-space R^2^ per trait under each split design. Open diamonds mark the published two-stage values, which used Split A’s design; Rdark25 has no published value and so carries no diamond. Both Rdark25 and Rday25 go negative under Split B. c) Leave-one-dataset-out skill for each held-out dataset, both tiers, ordered ascending by the baseline. The two-stage baseline is negative against its own null on four datasets (Montes et al. 2022, Heckmann et al. 2017, Albert et al. 2018, Rogers et al. 2017); the constrained model is negative on none. The pooled block of six datasets with fewer than 40 curves is shown for completeness and is excluded from every paired test, as marked. Tier 2 values are ensemble means over ten seeds.

### What the curve itself constrains

We next asked whether all four parameters are equally well defined by the gas-exchange measurement itself, independent of any spectral model, since two of the benchmark’s targets may be hard to identify from the curve before any spectrum is involved. Before using the differentiable FvCB decoder to answer that, we confirmed it reaches GSTI’s own fitted optimum rather than a systematically offset one. When fitted freely to each curve’s own measured points within each of GSTI’s four curve model classes, the median decoder-RMSE to reference-RMSE ratio is 0.9996–1.0000 (Table S2), and Rday25, the parameter the curve identifies best, is recovered at R^2^ = 0.985 across all 2983 curves with no fits at a bound (Fig. 3b). Any identifiability limit below is therefore a property of the data rather than of a mis-calibrated physics layer. We then stratified that free fit by curve model class to ask whether TPU25’s published R^2^ of 0.80 (Lamour *et al*., 2026) reflects a parameter the underlying data can identify. Within curves that sample the TPU-limited region (Ac_Aj_Ap, n = 805), fitted and curated TPU25 agree closely (R^2^ = 0.9993), with no fits at the parameter bound. Within curves that do not (Ac_Ap, n = 399), the agreement collapses and 80 of 399 fits sit at the upper bound of 200 µmol m^-2^ s^-1^ (Fig. 3a). The published TPU25 benchmark pools both classes. Progressively removing points from specific C_i_ regions and refitting shows that the same coverage logic governs every parameter. Thinning degrades TPU25 and Jmax25 most under high-C_i_ removal (correlation with the full-curve estimate 0.32 and 0.11, against 0.65 and 0.61 under uniform thinning), and Vcmax25 and Rday25 most under low-C_i_ removal (0.25 and 0.35, against 0.93 and 0.89; Fig. 3c). No FvCB parameter is uniformly easy or difficult to fit; each is identifiable only from a specific segment of C_i_ coverage. Taken together, whether a curve identifies TPU25 depends on which curves happen to sample that segment rather than on a general property of the parameter.

**Fig. 3.**
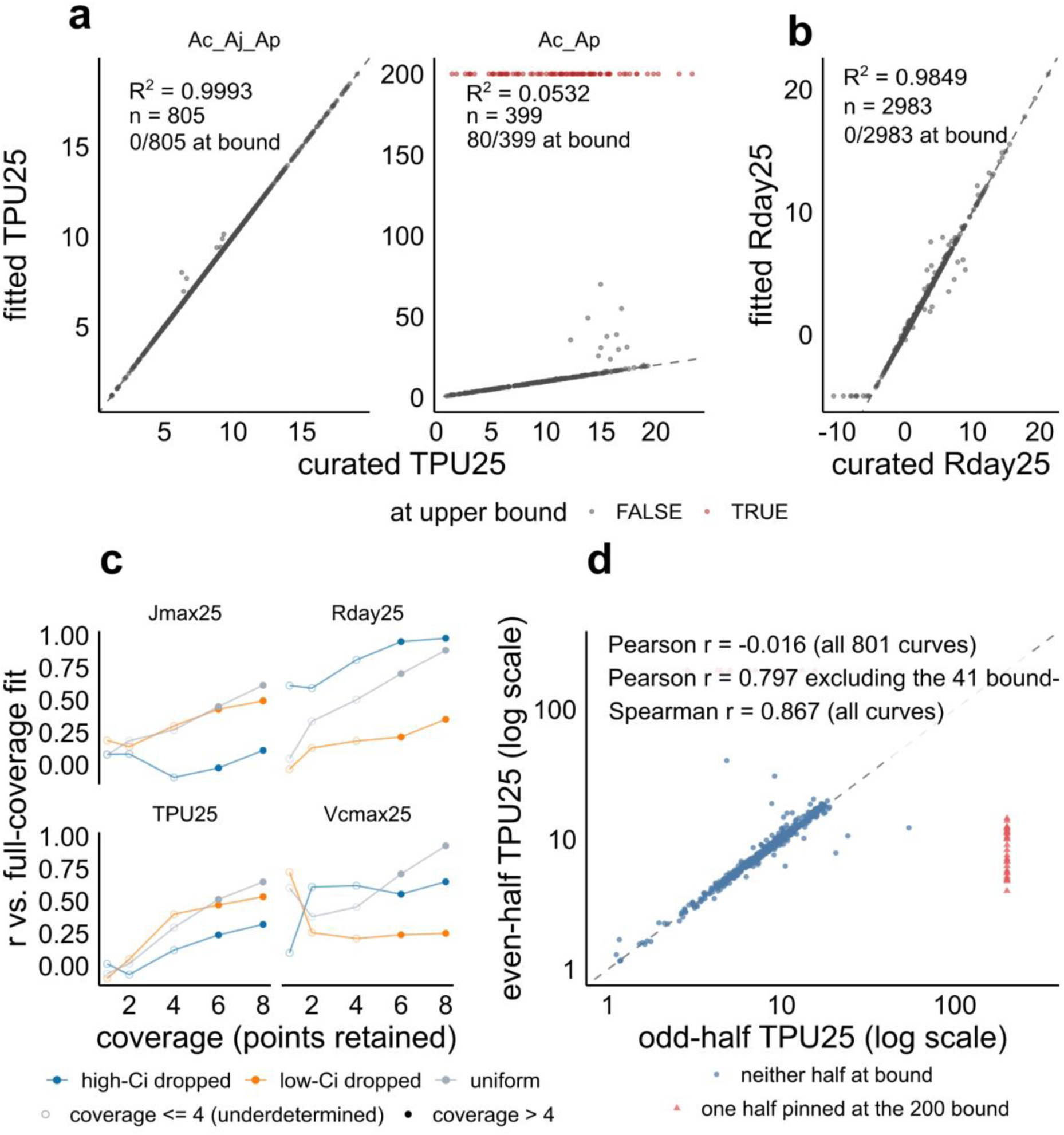
Gas exchange curve constraints. a) TPU25, Tier 0 fitted against curated, faceted by GSTI model class. Red points are fits sitting at the 200 µmol m^-2^ s^-1^ upper bound. Ac_Aj_Ap produces R^2^ = 0.9993, n= 805, 0/805 at bound. Ac_Ap produces R² = 0.0532, n = 399, 80/399 at bound. b) Rday25, Tier 0 fitted against curated (R^2^ = 0.9849, n = 2983, none at bound). c) Correlation with the full-coverage fit as points are removed under three thinning schemes, faceted by parameter. Hollow points mark coverage ≤ 4, at or below the four-point floor for a four-parameter fit. d) Split-half reliability of TPU25 within the Ac_Aj_Ap stratum. The odd-point fit against the even-point fit for all 801 curves, log axes. Red triangles are the 41 curves with one half pinned at the 200 bound.

### What the spectrum constrains

We next asked which of the curve-identifiable parameters are also identifiable from reflectance alone, and whether the model can recover them independently or not. We scored this under Split B and Split C rather than on the frozen 589-curve test set, which we established as a leaky design. Using our deep learning model, Vcmax25 and Jmax25 are recovered at r = 0.716 and 0.682 under Split B (Fig. 4a) and at 0.719 and 0.745 under Split C (Fig. 4b), against 0.815 and 0.803 on the leaky partition (Fig. S1a). However, TPU25 and Rday25 collapse into horizontal bands under both blocked designs, and the two failures have different causes. TPU25’s target is not reproducible across the database. Fitting each curve’s two halves independently gives a split-half reliability of 0.394, against 0.837 for Vcmax25 (Fig. 4c). It is reproducible within the identifiable stratum once 41 bound-pinned leverage curves are set aside (r = 0.797 against a naive −0.016; Fig. 3d) — of the 805 Ac_Aj_Ap curves, 801 converged in both half-fits and enter this comparison — but 45% of all curves place a half-fit at the bound (Fig. 4d). Curated Rday25, by contrast, is well determined (reliability 0.790, bounding any predictor near 0.94; Fig. 4c) and is still not recovered, at r = −0.149 under Split B and −0.134 under Split C, and at no curve coverage tested (Fig. S2b, Fig. S1b). The cause is architectural as ∂A/∂Rday = −1 everywhere in the decoder (Fig. S2a), so the assimilation loss barely responds to it (0.05% change in negative log-likelihood under perturbation, against 2.7% and 5.1% for Vcmax25 and Jmax25; Fig. S1d), while a directly supervised respiration head on the same encoder recovers Rdark25 (Fig. S2e; Methods S1). Neither of the two recovered parameters is recovered at full range or independently, and predictions are compressed toward the mean (Split B slopes 0.41 and 0.45, underpredicting the highest curated decile by 47% and 41%; Fig. S2d). Jmax25 retains 17% of its raw correlation once curated Vcmax25 is held constant (Fig. 4e), and the Jmax25:Vcmax25 ratio is not predicted above a training-median null (skill −0.087 and −0.063; Fig. 4f) despite the ratio’s own split-half reliability of 0.800. The two-stage baseline collapses the same way and more sharply, so this belongs to what leaf spectra carry rather than to the architecture. Taken together, once evaluation is blocked a spectrum constrains one compressed axis of photosynthetic capacity rather than four independent parameters.

**Fig. 4.**
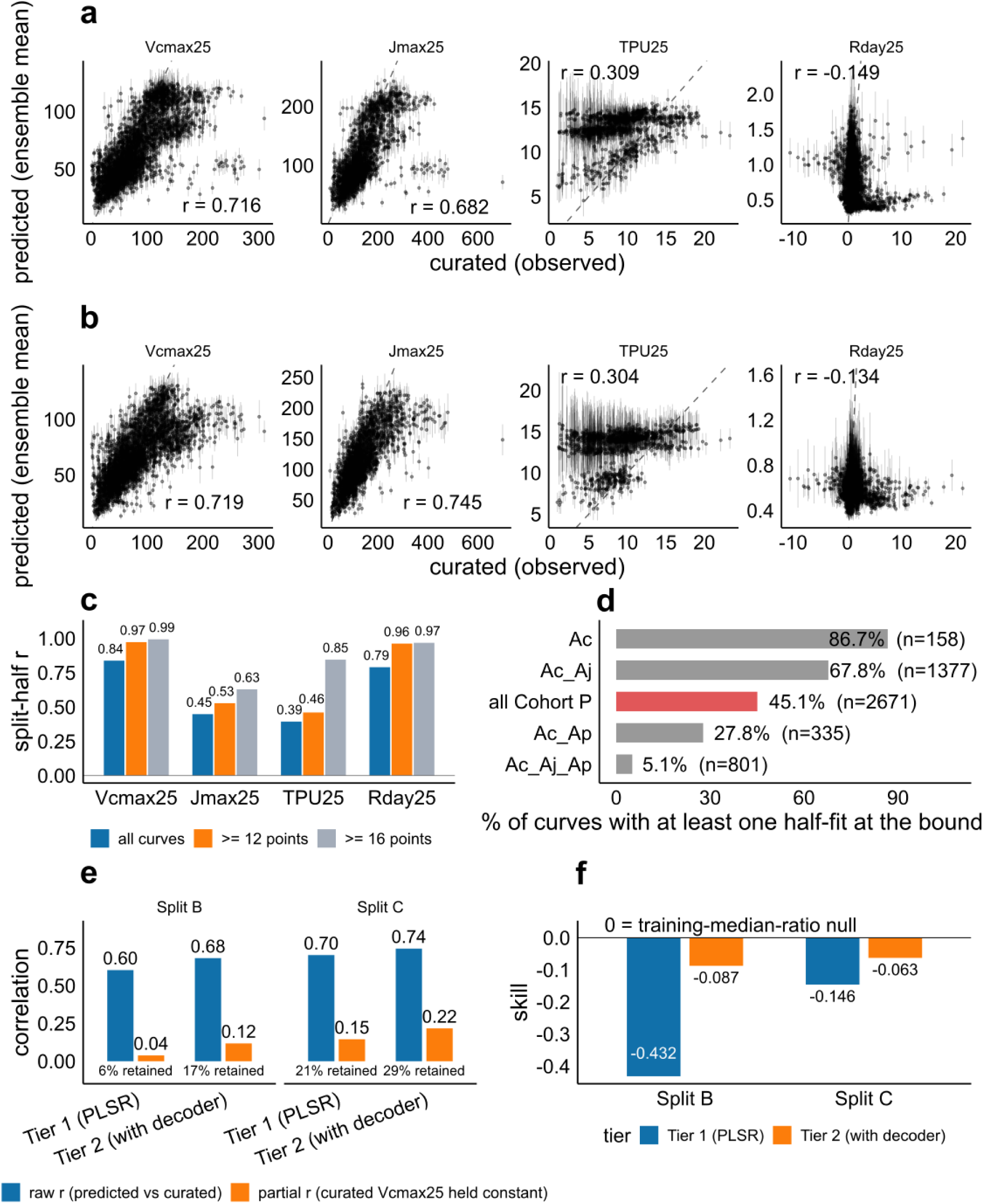
Spectrum constrains one axis of photosynthesis. a) Latent recovery under Split B (dataset-blocked) predicted against curated for each parameter, ensemble mean with error bars giving the seed standard deviation. The dashed line is 1:1. b) Latent recovery under Split C (species-blocked), same construction. c) Split-half reliability of each parameter at three coverage tiers where all curves, curves with at least 12 points, and curves with at least 16 points are shown. d) Fraction of curves with at least one half-fit at the TPU25 upper bound, by GSTI model class. Ac_Aj_Ap, the stratum in which TPU25 is nominally identifiable. e) Jmax25 recovery before and after holding curated Vcmax25 constant, both tiers, under each blocked design. f) Skill in predicting the Jmax25:Vcmax25 ratio against a training-median-ratio null.

### What the constraint costs and provides

Finally we asked what routing spectral predictions through fixed biochemistry costs relative to the two-stage baseline, what it provides, and whether the constraint itself is responsible for any advantage the architecture shows. Under the published within-dataset design, the constrained model (Tier 2, spectrum → four FvCB parameters → fixed decoder) underperforms the baseline (Tier 1, PLSR on individually fitted parameters); on the frozen 589-curve test partition, skill is 0.3642 ± 0.0323 against 0.4057. Under leave-one-dataset-out, Tier 2 is never negative relative to its own null across all 17 held-out datasets, whereas Tier 1 is negative on four. Additionally, Tier 2 exceeds Tier 1 on 16 of 17 (Wilcoxon signed-rank p = 0.0001, median paired improvement +0.168; Fig. 5a). Scored across all three designs, the advantage is specific to dataset-level transfer, where Tier 2 trails under the published design (0.409 against 0.439 by inner five-fold cross-validation over Split A’s training partition), but leads under dataset-blocked holdout (0.352 against 0.282) and is level under species-blocked holdout (0.261 against 0.259; Fig. 5b). We then eliminated the alternative explanations in turn. Encoder capacity does not account for the reversal. A grid over depth, width, kernel size, spectral binning and auxiliary loss weight spans 0.336–0.374 among the architecture-only arms, a spread of 0.037 that is narrower than the 0.066 gap to Tier 1, and the configuration ranks 7th of 10 (Fig. S3a,b; Table S1). Training-set size does not account for it either. Tier 2 trains on more curves per trait than the baseline can use (3860 against 1191–2895; Fig. S4a,b,c), but restricting it to the baseline’s own curves moves skill by 0.015 across a 2.5-fold span in training data (Fig. 5c, Fig. S4d). Preprocessing does not account for it either, since giving the baseline Tier 2’s own input representation changes its mean parameter-space correlation by −0.006 under Split B and −0.018 under Split C (Fig. S5a,b). We also tested the physics constraint, an otherwise identical encoder with the decoder removed, regressing directly on curated parameters, and the result is indistinguishable from Tier 2 under leave-one-dataset-out (0.411 against 0.417; Wilcoxon p = 1.00; Table 1). Taken together, the transfer advantage belongs to the model family rather than to the FvCB constraint, and the constraint is the instrument that made the parameter-by-parameter results measurable.

**Fig. 5:**
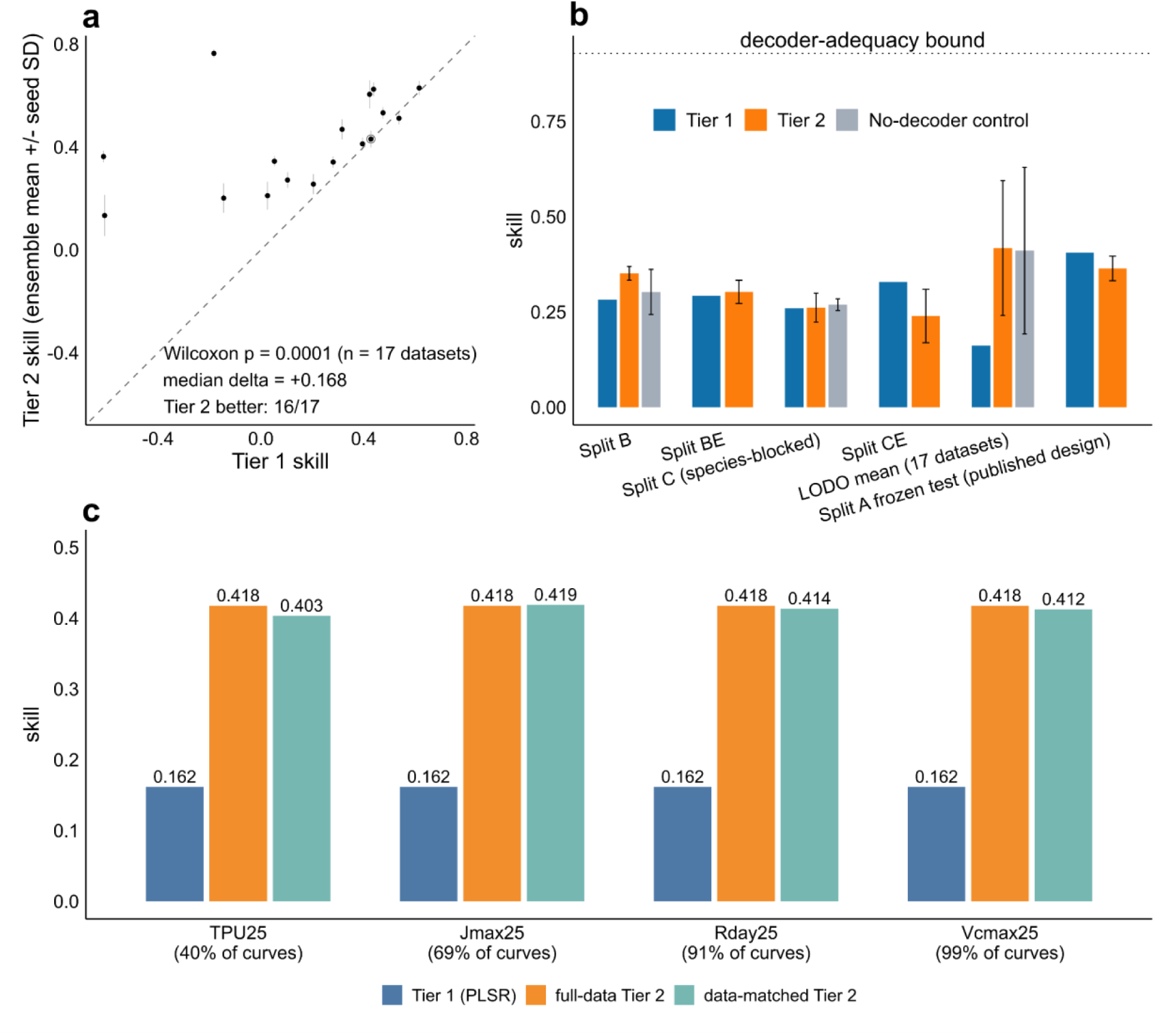
Physics constraints trade-off in information retrieval. a) Paired leave-one-dataset-out skill, 17 held-out datasets. Points above the 1:1 line are datasets where the constrained model beats the two-stage baseline; error bars give the seed standard deviation. Wilcoxon signed-rank p = 0.0001 at n = 17 datasets, median paired improvement +0.168, constrained model better on 16 of 17. The circled point (Cui et al. 2025) is the dataset that moved the count from 15/17 to 16/17 when the seed count rose from three to ten. b) Skill by fold definition, all blocking schemes, with the no-decoder control included. The control was run under Split B, Split C and LODO only. c) Data-matched ablation where the constrained model retrained on only the curves the two-stage baseline could use for each trait, against the full-data constrained model and the baseline, under LODO. The percentages denote the curves that were available for the prediction of each of the parameters.

**Table 1.** Skill of the constrained model, the no-decoder control and the two-stage baseline under each evaluation design.

| Condition | Split | Skill<br>(mean) | SD<br>(across<br>seeds) | SD<br>(across<br>datasets) | n<br>seeds | Compara<br>tor | p | n<br>compar<br>ed |
| --- | --- | --- | --- | --- | --- | --- | --- | --- |
| Tier 2 (with<br>decoder) | B | 0.3515 | — | — | 10 | — | — | — |
| No-decoder<br>control | B | 0.3026 | 0.0590 | — | 3 | Tier 2 | 0.288 | 15 |
| Tier 2 (with<br>decoder) | C | 0.2614 | — | — | 10 | — | — | — |
| No-decoder<br>control | C | 0.2692 | 0.0154 | — | 3 | Tier 2 | 0.473 | 15 |
| Tier 1<br>(PLSR<br>baseline) | LODO<br>(per<br>dataset) | 0.1616 | n/a | 0.3701 | — | — | — | — |
| Tier 2 (with<br>decoder) | LODO<br>(per<br>dataset) | 0.4176 | 0.0349 | 0.1767 | 10 | Tier 1 | $1.1 \times 10^{-4}$ | 17 |
| No-decoder<br>control | LODO<br>(per<br>dataset) | 0.4109 | 0.0519 | 0.2185 | 10 | Tier 1;<br>Tier 2 | $5.0 \times 10^{-4}$; 1.00 | 17 |

## Discussion

The GSTI benchmark evaluates interpolation rather than transfer, as revealed by its random split design within each contributing dataset (Lamour *et al*., 2026). Random splits overstate skill when data exhibit grouped structures, necessitating blocking of the structure that generates dependence (Roberts *et al*., 2017; Ploton *et al*., 2020). Here the relevant group is the dataset, not the species. Split B holds out entire contributing datasets, so a fold boundary coincides with every batch-level difference between them, including instrument, protocol, site and season. Split C holds out species and leaves dataset membership on both sides. Rdark25 separates the two designs. It recovers to R^2^ = 0.534 under species blocking, close to its within-dataset value of 0.585, and falls to −0.230 under dataset blocking. No other parameter diverges this way. This behavior indicates that Rdark25 depends more on the growth environment and the instrument producing a curve than on the species. Instrument, operator, and protocol are well-established latent grouping variables in high-throughput measurement (Leek *et al*., 2010), and leakage of this kind is not detected by ordinary validation (Kapoor & Narayanan, 2023; Stock *et al*., 2023). Leaf spectral models are already known to lose accuracy on new sites, seasons, and plant functional types (Burnett *et al*., 2021). This does not contradict GSTI’s original findings, which document its validation design and claim no more. What we quantify is the consequence of interpreting within-dataset interpolation scores as estimates of field performance. For this database, these differences are significant enough to reverse the sign of the result for two of five traits.

The transfer advantage under blocked evaluation is genuine, but it is not due to the physics constraint. An otherwise identical encoder without a decoder, regressing directly on curated parameters, transfers as well as the constrained model under leave-one-dataset-out. Analysis revealed 0.411 against 0.417, with a mean paired difference of −0.007 across the 17 held-out datasets and a bootstrap interval of [−0.053, +0.039] (Table 1). The two one-sided tests do not clear our pre-specified equivalence margin of ±0.033 (p = 0.060), indicating that any true advantage of the constrained model is too small to detect at the replication level rather than demonstrably absent. Encoder capacity, training-set size, and spectral preprocessing were each tested directly and excluded. What remains is the model family. A convolutional encoder trained on whole curves transfers across datasets where per-trait linear regression on fitted parameters does not, and the baseline’s skill varies more than twice as much across held-out datasets compared to the constrained model’s (SD 0.370 against 0.177). Hybrid architectures that embed mechanistic equations inside a network are now widely advocated (Reichstein *et al*., 2019; Aboelyazeed *et al*., 2023), and are usually credited on the strength of a capacity sweep. However, a capacity sweep cannot separate an architecture that helps due to its physical structure from one that benefits because of the function class it belongs to, as every arm retains the constraint. The no-constraint arm serves as a control that separates them, and comparable audits show that reported advantages often do not survive stronger controls (McGreivy & Hakim, 2024). Instead, the decoder provides measurement. Perturbing one parameter while holding others fixed, or conditioning Jmax25 recovery on Vcmax25, is a defined operation only because the parameters enter an explicit physiological model (Raue *et al*., 2009). A purely predictive network returns four numbers without a way to determine which of them the spectrum constrains.

Two of the four parameters fail for different reasons, and neither reason is that spectra carry no information about them. Curated TPU25 is not reproducible enough to serve as a pooled benchmark target. Its split-half reliability is 0.394 across the database and only 0.460 on curves with at least twelve points, with 45% of curves placing a half-fit at the fitter’s upper bound of 200 µmol m^-2^ s^-1^, roughly twenty times the median curated value. Bound-hitting fits are a documented property of A-C_i_ fitting software (Lochocki *et al*., 2025), and the choice of fitting tool alone spreads fitted parameters widely across the same curves (Wei *et al*., 2026). Within the stratum where a curve reaches the TPU-limited region (Gu *et al*., 2010; Sharkey, 2019) the independent halves agree, giving r = 0.797 once the 41 bound-pinned curves are set aside, but most curves never sample that region, and the spectral model fails inside the stratum in any case (R^2^ = −1.18, against −0.41 on curves that cannot identify TPU25 at all). TPU25 is identifiable in principle and not from this measurement design as demonstrated in other analogous physiology modeling domains (Riedlinger *et al*., 2015). Rday25 fails on the opposite terms. Its curated values are well determined, with a split-half reliability of 0.790 rising to 0.962 at twelve points or more, but the loss barely responds to it. Perturbing each predicted parameter by a tenth of its interquartile range changes the negative log-likelihood by 2.7% for Vcmax25 and 5.1% for Jmax25, and by 0.05% for Rday25. A second respiration head on the same encoder, supervised directly, recovers dark respiration where the decoder-routed head does not (r = 0.324 against −0.007 under dataset blocking, on the paired subset of curves carrying both respiration measurements; the decoder-routed head’s Split B recovery of Rday25 across all held-out curves is r = −0.149). Rdark and Rday are physiologically distinct quantities (Abadie *et al*., 2024; Yin & Amthor, 2024), so this is an internal control rather than a like-for-like substitute. A parameter that enters an assimilation-space loss as an additive offset is not separately identifiable through that loss, and loss surfaces of this shape are a recognised failure of the optimisation problem rather than of model capacity (Krishnapriyan *et al*., 2021).

Predictive skill and parameter identifiability are separate here, with the disagreement between them being most consequential for downstream use. Replacing each predicted parameter in turn with its cohort median shows that assimilation-space skill is carried mostly by Jmax25 (skill falls by 0.201), with Rday25 contributing almost nothing (0.009), while the identifiability ordering is close to inverted, since Jmax25 retains only 17% of its raw recovery once curated Vcmax25 is held constant. A model can improve the assimilation fit through a parameter it does not identify, as the fit requires only the combination of parameters that the spectrum constrains. Assimilation skill was our primary metric and we use it because it is the only currency in which every curve can be scored under transfer, and the parameter-space and partial correlations carry the identifiability claims. Recovery is also compressed toward the mean, with Split B slopes of 0.41 for Vcmax25 and 0.45 for Jmax25, and the highest curated decile underpredicted by 47% and 41%, respectively. This attenuation at sparsely sampled extremes is general to spectral trait models (Kothari *et al*., 2023). The model does not predict Jmax25:Vcmax25 above a training-median null (−0.087 and −0.063), although the ratio’s own split-half reliability of 0.800 leaves ample headroom. Its four outputs load onto two latent dimensions rather than four (99.8% of variance in the first two), with predicted Vcmax25 and predicted Jmax25 correlating at r = 0.995 against a curated correlation of 0.942 (Fig. S1e). This is the expected outcome if spectra track photosynthetic capacity through covarying leaf constituents rather than through separate signals for each parameter (Kothari & Schweiger, 2022). Earth system models need carboxylation and electron transport capacity to vary independently, and the ratio between them varies with growth temperature, climate and acclimation state rather than holding at a constant value (Kattge & Knorr, 2007; Rogers *et al*., 2017; Walker *et al*., 2017; Smith *et al*., 2019). Two limits bound these conclusions. Species blocking removes conspecific overlap but not phylogenetic structure, and leaf spectra carry phylogenetic signals (Meireles *et al*., 2020), so our species-blocked recovery is an upper bound on transfer to a phylogenetically novel leaf. We also cannot yet say why four of the seventeen datasets fail under leave-one-dataset-out. Spectral distance from the training distribution is the obvious candidate and is testable with existing area-of-applicability methods (Meyer & Pebesma, 2022).

A benchmark number is only as informative as the design that produced it. A design not checked against the non-independence structure of its own data can be wrong in either direction, understating transfer or overstating identifiability, and nothing in the number indicates which has occurred. Each result that initially appeared inconsistent here resolved once interpreted as a consequence of what a given design holds fixed, the divergence of Rdark25 between blocking schemes, the reversal between the published split and leave-one-dataset-out methods, and the confounded TPU25 score. What remains is a narrower claim about leaf reflectance than the field has been working with, and a sharper one. A spectrum resolves one axis of photosynthetic capacity. The balance between carboxylation and electron transport, which is the quantity Earth system models most need from these data, is not on that axis, and recovering it will require information that a leaf reflectance spectrum does not carry.

## Conclusion

The broader value of this work lies in the approach it provides in understanding what a spectral measurement can support. We show spectral reflectance constrains photosynthetic capacity along one dimension rather than independently resolving its components. This kind of limitation is exactly the type of characterization that is needed for future airborne and satellite imaging spectroscopy missions to scope out their utility. Similarly, whenever a data-driven model is coupled with a process representation, whether predicting gross primary productivity from canopy reflectance or inferring biochemical parameters for land-surface models, the question of what observations actually constrain and whether reported accuracy numbers reflect transfer or interpolation becomes critical. This question can be addressed using grouped evaluation and a differentiable forward model. Treating this problem as a fundamental part of model development, is essential for spectral and hybrid approaches to gain the confidence needed to inform carbon-cycle projections at the scales that motivate them.

## Author contribution

Conceptualization: RR; Data curation: RR; Formal analysis: RR; Funding acquisition: JM and TM; Investigation: RR; Methodology: RR; Supervision: JM and TM; Visualization: RR; Writing - Original Draft: RR; Writing - Review & Editing: RR, JM and TM

## Data Availability

All the raw data is available at https://github.com/plantphys/gsti. The data and codes used in this study is available at https://github.com/rishavray/dl_spectra_photosynthesis.

## Acknowledgements

This research was funded by the National Science Foundation (DEB-2129589 to JM, TM) and The National Institute of Food and Agriculture, United States Department of Agriculture (USDA-NIFA) award CA-D-PLB-2795-H to JM.

## Supplementary Figures

**Fig. S1:**
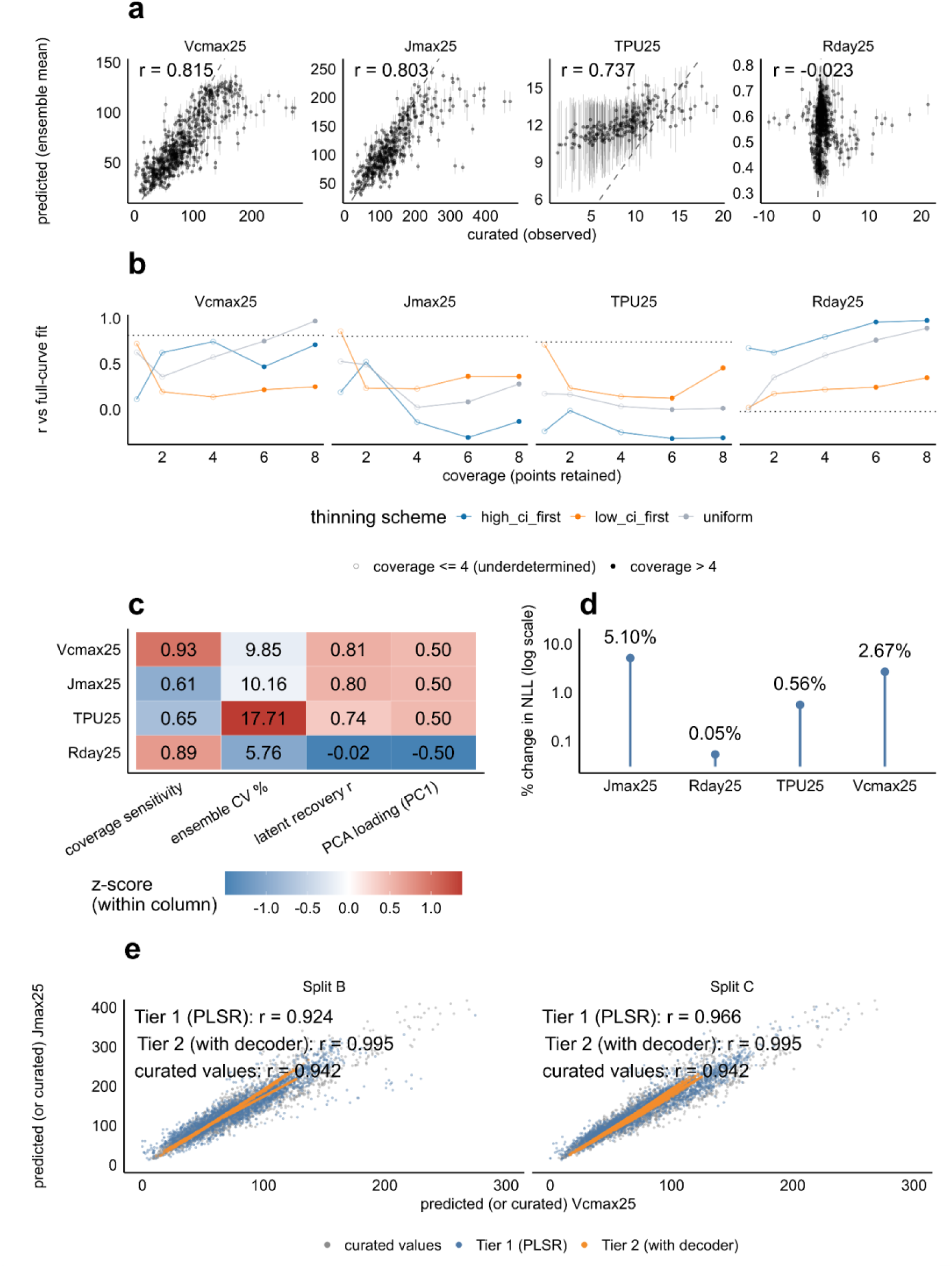
Recovery under the published design, and diagnostics that bound every correlation. (a) Latent recovery under Split A, the published within-dataset design where ensemble-mean is predicted against curated values for each of the four FvCB parameters, with the Pearson correlation annotated (Vcmax25 r = 0.815, Jmax25 0.803, TPU25 0.737, Rday25 −0.023). (b) Curve-only information by coverage. Tier 0 correlation with the full-curve fit as points are thinned from each A–C_i curve (8 to 1 retained), under three removal schemes (uniform, high-C_i-dropped, low-C_i-dropped), with the level a spectrum alone achieves (Tier 2) marked as a reference line per parameter. Filled symbols are coverage > 4; open symbols are the underdetermined regime (≤ 4 points). (c) Agreement of four independent diagnostics on the parameter ranking, shown as z-scores computed within each column. Coverage sensitivity, cross-seed ensemble coefficient of variation, latent recovery r, and loading on the first principal component of the encoder’s four outputs. (d) Loss-surface response to each parameter. The change in held-out assimilation negative log-likelihood when each predicted parameter in turn is perturbed by 10% of its own interquartile range across Cohort P (2.7% for Vcmax25, 5.1% for Jmax25, ≈ 0.05% for Rday25). The loss is nearly flat in Rday25, which is the basis for the architectural reading of its failure. (e) Predicted Vcmax25 against predicted Jmax25 for both tiers under both blocked designs, with the curated Vcmax25–Jmax25 relationship overlaid for reference. The two predicted capacities correlate at r = 0.995 for the constrained model (both splits) and 0.924 (Split B) / 0.966 (Split C) for the baseline, against r = 0.942 between the curated values.

**Fig. S2:**
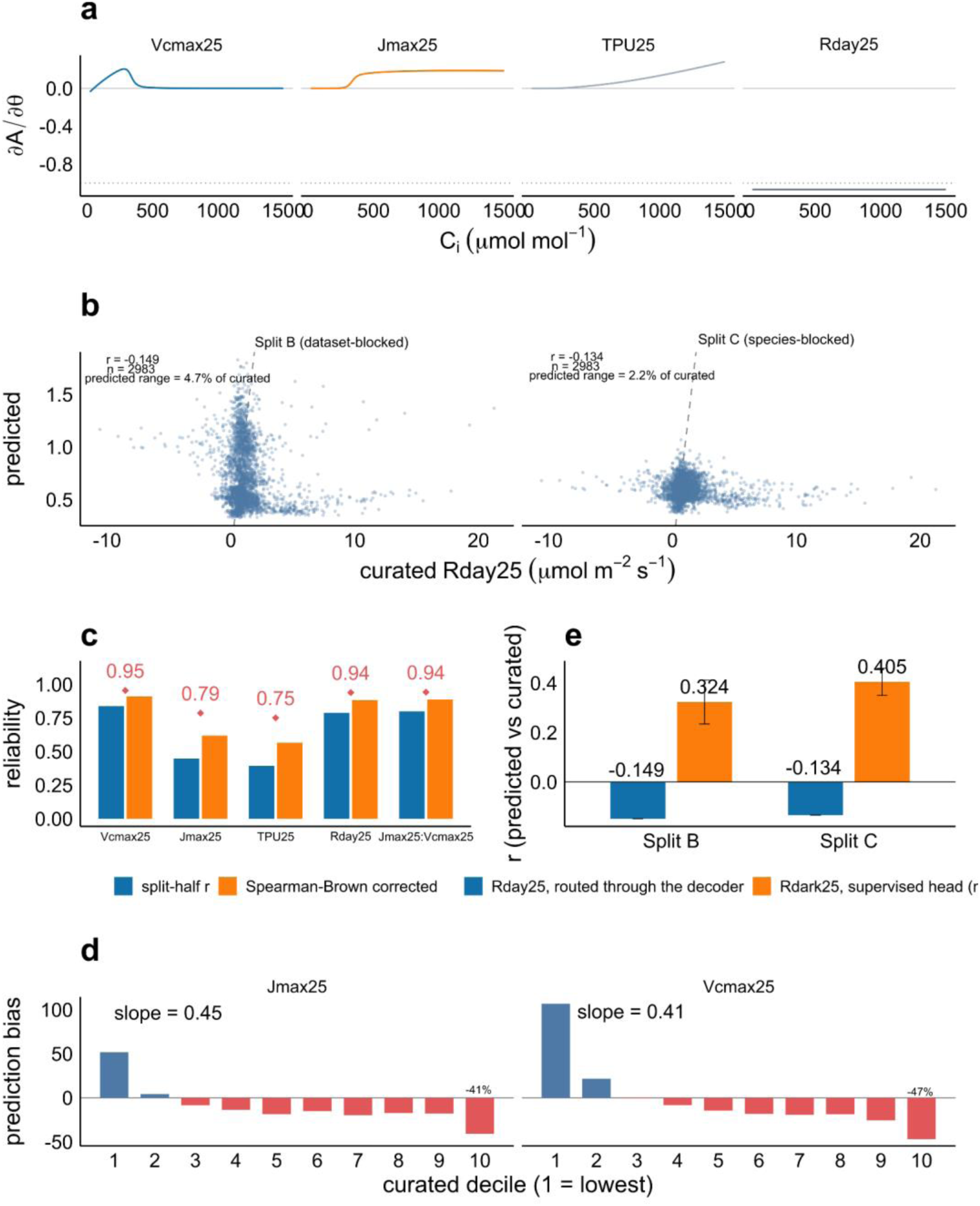
what the spectrum constrains, methodological details. (a) Analytic sensitivity of decoded net assimilation to each FvCB parameter, ∂A/∂θ, computed by automatic differentiation through the fixed decoder at Cohort P’s median parameter vector and plotted point-by-point along the C_i axis rather than on a summed loss. The Vcmax25, Jmax25 and TPU25 derivatives are localized to specific C_i regions, whereas ∂A/∂Rday25 = −1 everywhere. Respiration enters as a uniform additive offset carrying no positional information for the loss to exploit. (b) Rday25 predicted against curated under Split B (dataset-blocked) and Split C (species-blocked), ensemble means over ten seeds. Predictions collapse into a horizontal band spanning 4.7% (Split B) and 2.2% (Split C) of the curated range (r = −0.149 and −0.134, n = 2983), the signature of a head emitting approximately one value per dataset. (c) Split-half reliability of the four FvCB parameters and of the Jmax25:Vcmax25 ratio, from fitting each curve’s odd- and even-indexed points independently. Raw split-half r and the Spearman–Brown-corrected value, with the red diamond marking √r_SB, the ceiling any predictor could reach against that curated target. (d) Range compression under Split B. Mean signed prediction bias within each curated decile, expressed as a percentage of that decile’s own curated mean, with the predicted-on-curated regression slope annotated (Jmax25 slope 0.45, Vcmax25 0.41). (e) Controlled architectural comparison isolating target routing from spectral information. The same encoder and the same splits, differing only in whether respiration is reached through the decoder (Rday25) or through a directly supervised head (Rdark25).

**Fig. S3:**
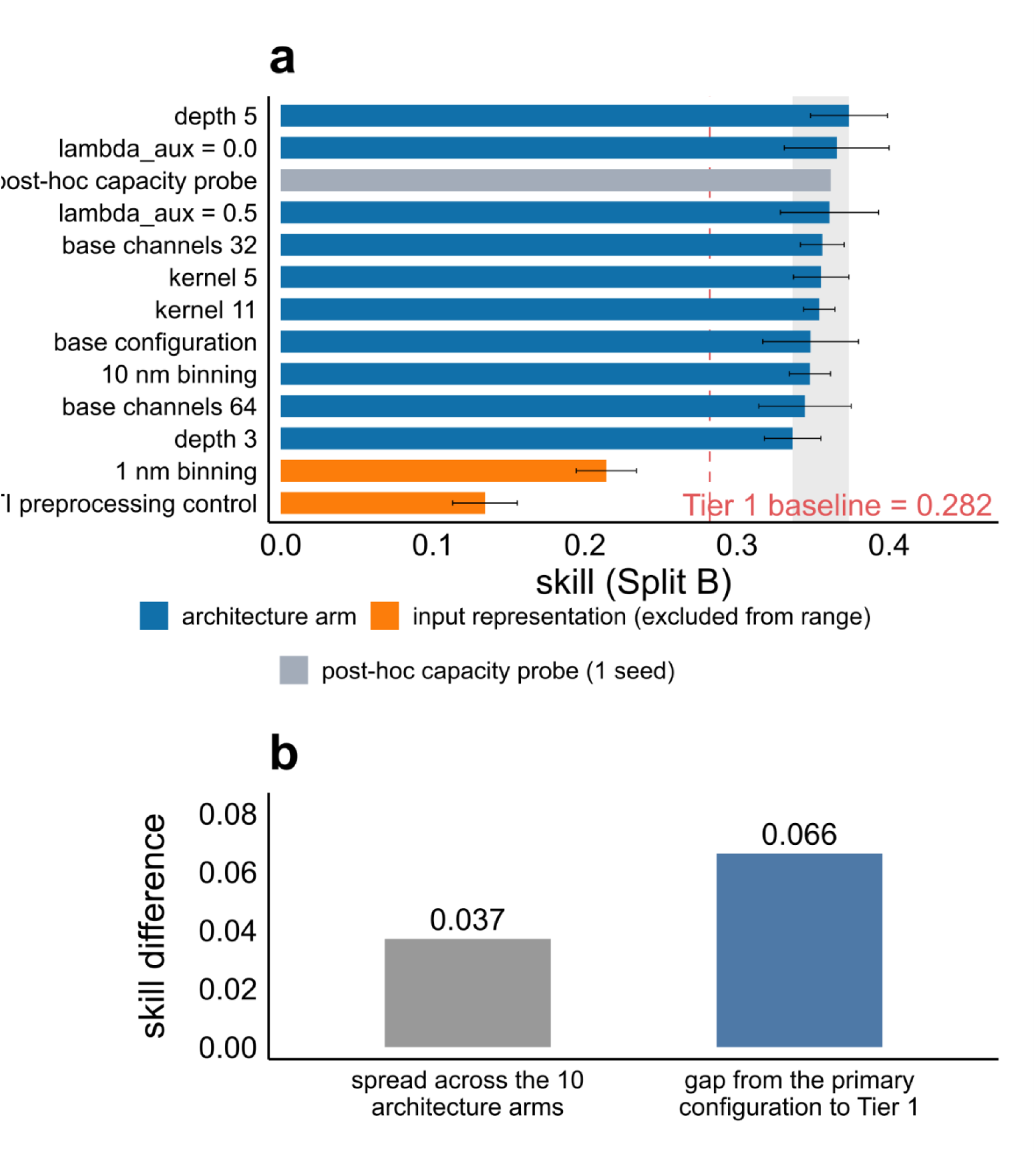
encoder capacity does not explain the transfer reversal. (a) Split B skill for every arm of the grid, ordered ascending, with seed standard deviations at three seeds per arm. Blue bars are the ten architecture-only arms (depth, channel width, kernel size, auxiliary loss weight, 10 nm binning), whose span is shaded. The dashed red line is the Tier 1 baseline at 0.282. Orange bars are the two arms that change the input representation rather than encoder capacity (1 nm binning at 0.214 - GSTI-preprocessing control at 0.134) and are excluded from the architecture range (b) The spread across the ten architecture arms (0.037) against the gap from the primary configuration to Tier 1 (0.066). The full range achievable by architecture choice is smaller than the gap it would need to explain, so capacity and tuning do not account for the reversal.

**Fig. S4:**
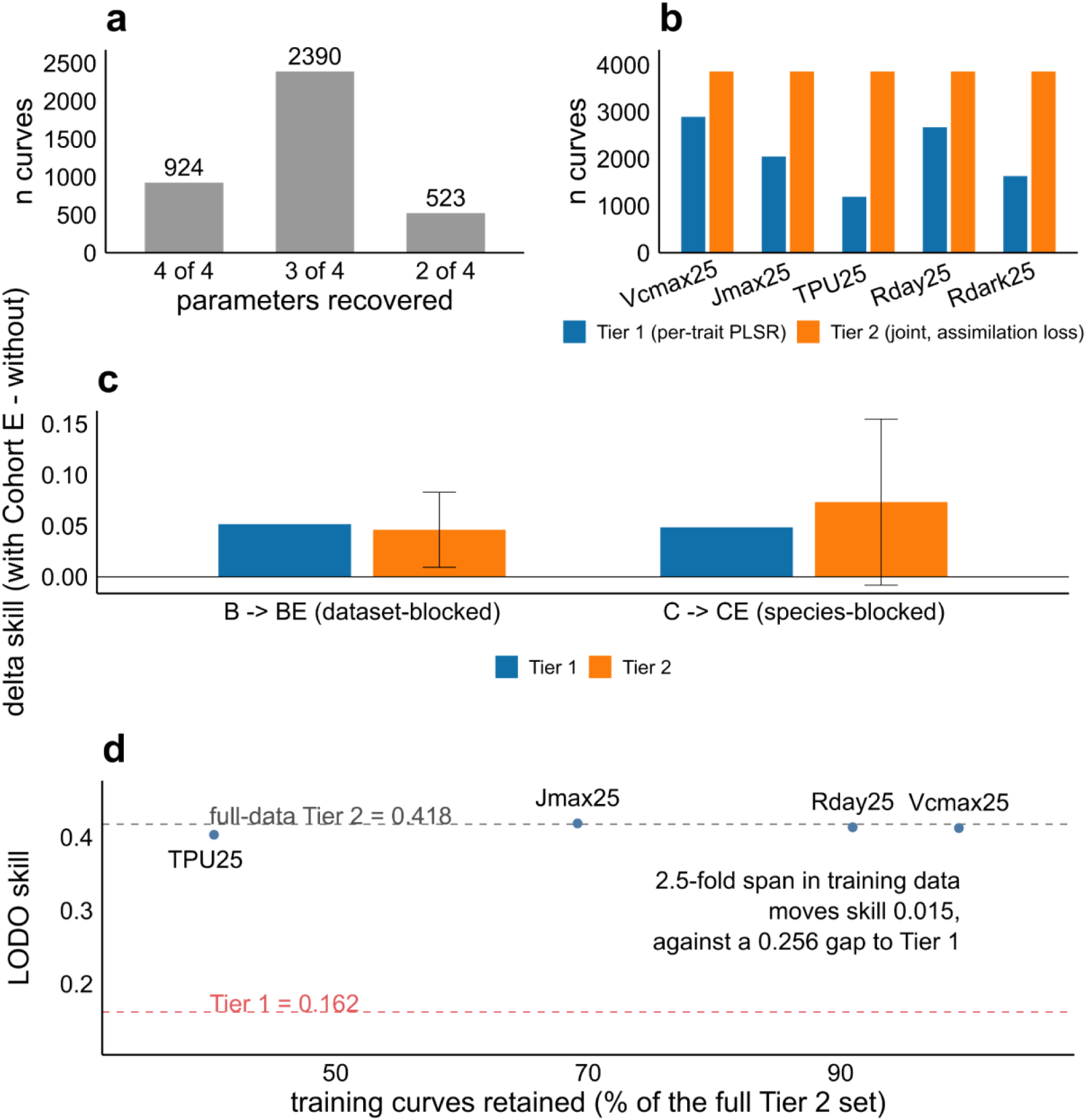
Training-set size does not explain the reversal. (a) Cohort P curves by the number of FvCB parameters GSTI’s pipeline recovered, database-wide. 924 curves with all four, 2390 with three, 523 with two. Most curves are not fully fit, which is what creates the difference in available training data between the two methods. (b) Training curves available per trait, baseline against constrained model. The constrained model uses all 3860 curves for every trait whereas the baseline can use only curves for which that trait was individually fitted (1191–2895 depending on trait). (c) Skill gain from adding Cohort E curves to the training portion, by tier and blocking scheme (B → BE, C → CE), with seed standard deviations where an ensemble exists. (d) Data-matched ablation where the constrained model retrained from scratch on only the curves the baseline could use for each trait, one restricted model per trait, scored under leave-one-dataset-out against the full-data constrained model (grey dashed, 0.418) and the baseline (red dashed, 0.162).

**Fig. S5:**
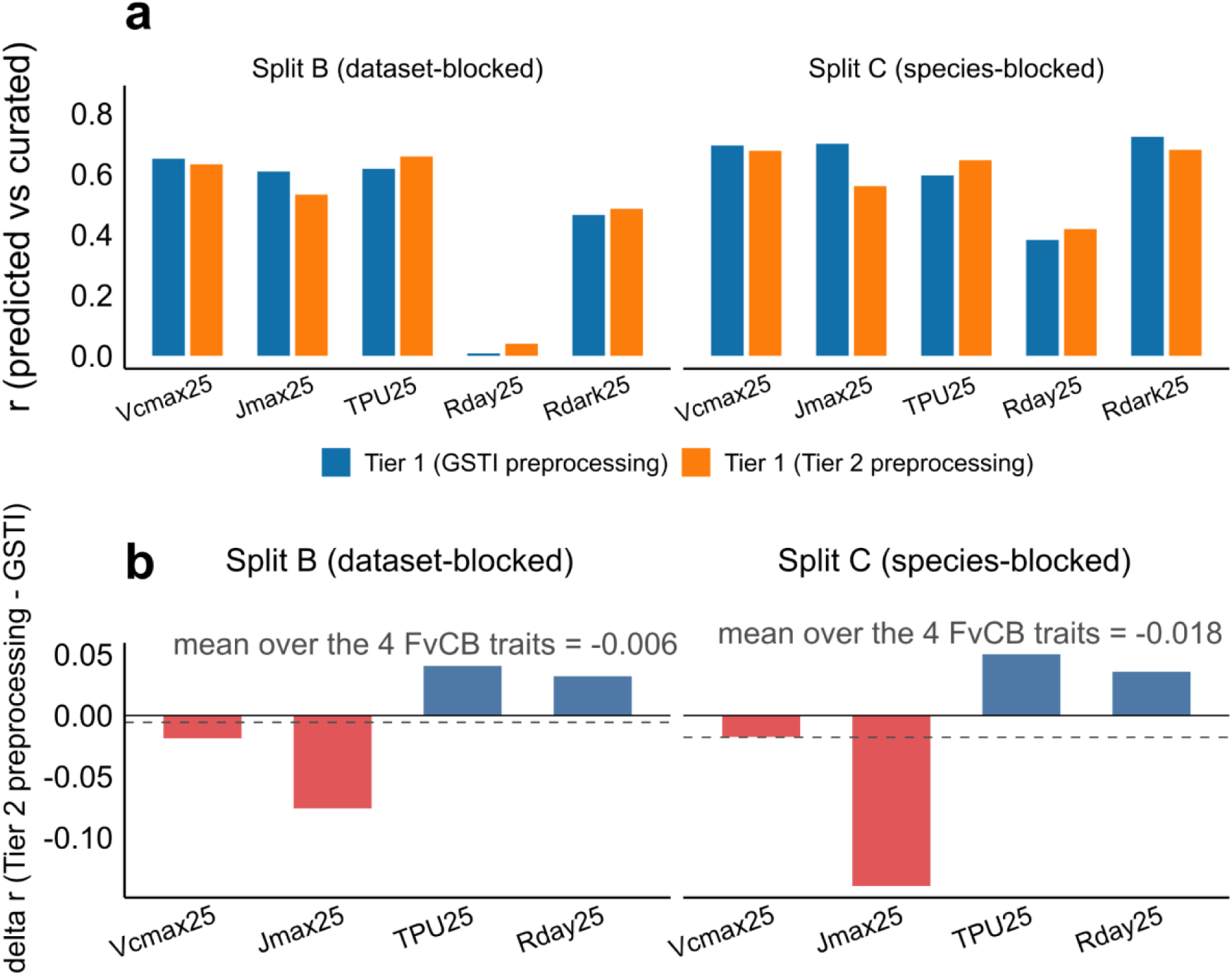
preprocessing does not explain the transfer reversal. (a) Parameter-space correlation (predicted against curated) for the baseline’s own PLSR algorithm, unchanged including its component-selection step, run on the constrained model’s preprocessing (5 nm binning, per-observation vector normalization, added first-derivative channel) in place of GSTI’s, under Split B and Split C, per parameter. Component counts were re-selected under each preprocessing. (b) The same result as per-trait differences (Tier 2 preprocessing minus GSTI preprocessing), with the mean over the four FvCB traits marked (−0.006 under Split B, −0.018 under Split C). The swap gains on TPU25 and Rday25 and loses on Jmax25, netting near zero, so preprocessing does not close the transfer gap in either direction.

**Table S1.** Sensitivity grid. Arm-by-arm Split B skill (mean ± seed standard deviation, three seeds each) for the post-hoc sensitivity grid, with the Tier 1 baseline for reference. Each row varies one factor from the primary configuration (depth, channel width, kernel size, spectral binning, or auxiliary loss weight λ_aux). The “in range” column marks which arms count toward the architecture-only span: the two arms that alter the input representation (1 nm binning; GSTI-preprocessing control) and the single-seed post-hoc capacity probe are excluded. The grid was run only after the primary configuration was locked and was not used to select it.

| Configuration | Depth | Width | Kernel | Bin (nm) | $\lambda_{\text{aux}}$ | Split | Skill (mean ± seed SD) | In range statistic |
| --- | --- | --- | --- | --- | --- | --- | --- | --- |
| Tier 1 (PLSR baseline) | — | — | — | — | — | B | 0.282 | No |
| Baseline configuration | 4 | 128 | 7 | 5 | 0.1 | B | 0.348 ± 0.031 (n = 3) | Yes |
| Depth 5 | 5 | 128 | 7 | 5 | 0.1 | B | 0.374 ± 0.025 (n = 3) | Yes |
| Depth 3 | 3 | 128 | 7 | 5 | 0.1 | B | 0.336 ± 0.019 (n = 3) | Yes |
| Kernel 5 | 4 | 128 | 5 | 5 | 0.1 | B | 0.355 ± 0.018 (n = 3) | Yes |
| Kernel 11 | 4 | 128 | 11 | 5 | 0.1 | B | 0.354 ± 0.010 (n = 3) | Yes |
| Base channels 32 | 4 | 32 | 7 | 5 | 0.1 | B | 0.356 ± 0.014 (n = 3) | Yes |
| Base channels 64 | 4 | 64 | 7 | 5 | 0.1 | B | 0.345 ± 0.030 (n = 3) | Yes |
| $\lambda_{\text{aux}} = 0.0$ | 4 | 128 | 7 | 5 | 0 | B | 0.366 ± 0.034 (n = 3) | Yes |
| $\lambda_{\text{aux}} = 0.5$ | 4 | 128 | 7 | 5 | 0.5 | B | 0.361 ± 0.032 (n = 3) | Yes |
| 10 nm binning | 4 | 128 | 7 | 10 | 0.1 | B | 0.348 ± 0.013 (n = 3) | Yes |
| 1 nm binning (excluded from range) | 4 | 128 | 7 | 1 | 0.1 | B | 0.214 ± 0.020 (n = 3) | No |
| GSTI<br>preprocess<br>ing control | 4 | 128 | 7 | 1 | 0.1 | B | $0.134 \pm 0.021$ (n = 3) | No |
| Post-hoc<br>capacity<br>probe | 6 | 256 | 7 | 5 | 0.1 | B | $0.362$ (n = 1) | No |

**Table S2.** Decoder validation. Agreement between Tier 0 (the decoder’s own free fit to each curve’s measured points) and GSTI’s curated values, establishing that the fixed decoder reaches GSTI’s fitting optimum rather than a systematically offset one.

| Output quantity | Slope | R<sup>2</sup> | RMSE | RMSE/$\sigma$ | n |
| --- | --- | --- | --- | --- | --- |
| V<sub>c</sub>max25 | 1.0041 | 0.9979 | 2.2110 | – | 2983 |
| J<sub>max</sub>25 | 1.0069 | 0.9985 | 2.9709 | – | 2063 |
| TPU25 | 2.9509 | 0.0531 | 48.7379 | – | 1204 |
| R<sub>day</sub>25 | 0.9781 | 0.9849 | 0.2366 | – | 2983 |
| A<sub>net</sub> (Ac model class) | – | – | 0.4756 | 1.0000 | 169 |
| A<sub>net</sub> (Ac_Aj model class) | – | – | 0.3712 | 0.9999 | 1610 |
| A<sub>net</sub> (Ac_Aj_Ap model class) | – | – | 0.3201 | 1.0000 | 805 |
| A<sub>net</sub> (Ac_Ap model class) | – | – | 0.4856 | 0.9996 | 399 |

## References

Abadie C, Lalande J, Dourmap C, Limami AM, Tcherkez G. 2024. Leaf day respiration involves multiple carbon sources and depends on previous dark metabolism. Plant, Cell & Environment 47: 2146–2162.

Aboelyazeed D, Xu C, Gu L, Luo X, Liu J, Lawson K, Shen C. 2025. Inferring Plant Acclimation and Improving Model Generalizability With Differentiable Physics-Informed Machine Learning of Photosynthesis. Journal of Geophysical Research: Biogeosciences 130: e2024JG008552.

Aboelyazeed D, Xu C, Hoffman FM, Liu J, Jones AW, Rackauckas C, Lawson K, Shen C. 2023. A differentiable, physics-informed ecosystem modeling and learning framework for large-scale inverse problems: demonstration with photosynthesis simulations. Biogeosciences 20: 2671–2692.

Audebert N, Le Saux B, Lefevre S. 2019. Deep Learning for Classification of Hyperspectral Data: A Comparative Review. IEEE Geoscience and Remote Sensing Magazine 7: 159–173.

Beer C, Reichstein M, Tomelleri E, Ciais P, Jung M, Carvalhais N, Rödenbeck C, Arain MA, Baldocchi D, Bonan GB, et al. 2010. Terrestrial Gross Carbon Dioxide Uptake: Global Distribution and Covariation with Climate. Science 329: 834–838.

Burnett AC, Anderson J, Davidson KJ, Ely KS, Lamour J, Li Q, Morrison BD, Yang D, Rogers A, Serbin SP. 2021. A best-practice guide to predicting plant traits from leaf-level hyperspectral data using partial least squares regression. Journal of Experimental Botany 72: 6175–6189.

Busch FA, Ainsworth EA, Amtmann A, Cavanagh AP, Driever SM, Ferguson JN, Kromdijk J, Lawson T, Leakey ADB, Matthews JSA, et al. 2024. A guide to photosynthetic gas exchange measurements: Fundamental principles, best practice and potential pitfalls. Plant, Cell & Environment 47: 3344–3364.

Cui C, Fearn T. 2018. Modern practical convolutional neural networks for multivariate regression: Applications to NIR calibration. Chemometrics and Intelligent Laboratory Systems 182: 9–20.

Cui B, Mariani R, Cathline KA, Robertson G, Martin AR. 2025. Reflectance spectroscopy predicts leaf functional traits across wine grape cultivars. PLANTS, PEOPLE, PLANET 7: 1524–1537.

Deng X, Hu X, Shi L, Su C, Li J, Du S, Li S. 2025. Deep learning-enabled exploration of global spectral features for photosynthetic capacity estimation. Frontiers in Plant Science 15.

ElGhawi R, Kraft B, Reimers C, Reichstein M, Körner M, Gentine P, Winkler AJ. 2023. Hybrid modeling of evapotranspiration: inferring stomatal and aerodynamic resistances using combined physics-based and machine learning. Environmental Research Letters 18: 034039.

Fang J, Gentine P. 2024. Exploring Optimal Complexity for Water Stress Representation in Terrestrial Carbon Models: A Hybrid-Machine Learning Model Approach. Journal of Advances in Modeling Earth Systems 16: e2024MS004308.

Farquhar GD, von Caemmerer S, Berry JA. 1980. A biochemical model of photosynthetic CO2 assimilation in leaves of C3 species. Planta 149: 78–90.

Fu P, Meacham-Hensold K, Guan K, Bernacchi CJ. 2019. Hyperspectral Leaf Reflectance as Proxy for Photosynthetic Capacities: An Ensemble Approach Based on Multiple Machine Learning Algorithms. Frontiers in Plant Science 10.

Furbank RT, Silva-Perez V, Evans JR, Condon AG, Estavillo GM, He W, Newman S, Poiré R, Hall A, He Z. 2021. Wheat physiology predictor: predicting physiological traits in wheat from hyperspectral reflectance measurements using deep learning. Plant Methods 17: 108.

Gu L, Pallardy SG, Tu K, Law BE, Wullschleger SD. 2010. Reliable estimation of biochemical parameters from C3 leaf photosynthesis–intercellular carbon dioxide response curves. Plant, Cell & Environment 33: 1852–1874.

Hong D, Han Z, Yao J, Gao L, Zhang B, Plaza A, Chanussot J. 2022. SpectralFormer: Rethinking Hyperspectral Image Classification With Transformers. IEEE Transactions on Geoscience and Remote Sensing 60: 1–15.

Hong D, Zhang B, Li X, Li Y, Li C, Yao J, Yokoya N, Li H, Ghamisi P, Jia X, et al. 2024. SpectralGPT: Spectral Remote Sensing Foundation Model. IEEE Transactions on Pattern Analysis and Machine Intelligence 46: 5227–5244.

Huang Y, Liu N, Chen W, Tan W, Deng Y, Wang Z, Chlus A, Shen J, Townsend PA. 2026. Enhancing transferability of foliar trait retrieval models: A comparative analysis of transfer learning strategies and domain shift characterization. Remote Sensing of Environment 337: 115336.

Jacquemoud S, Baret F. 1990. PROSPECT: A model of leaf optical properties spectra. Remote Sensing of Environment 34: 75–91.

Kapoor S, Narayanan A. 2023. Leakage and the reproducibility crisis in machine-learning-based science. Patterns 4: 100804.

Kattge J, Knorr W. 2007. Temperature acclimation in a biochemical model of photosynthesis: a reanalysis of data from 36 species. Plant, Cell & Environment 30: 1176– 1190.

Kothari S, Beauchamp-Rioux R, Blanchard F, Crofts AL, Girard A, Guilbeault-Mayers X, Hacker PW, Pardo J, Schweiger AK, Demers-Thibeault S, et al. 2023. Predicting leaf traits across functional groups using reflectance spectroscopy. New Phytologist 238: 549–566.

Kothari S, Schweiger AK. 2022. Plant spectra as integrative measures of plant phenotypes. Journal of Ecology 110: 2536–2554.

Krishnapriyan AS, Gholami A, Zhe S, Kirby RM, Mahoney MW. 2021. Characterizing possible failure modes in physics-informed neural networks. In: NIPS ‘21. Proceedings of the 35th International Conference on Neural Information Processing Systems. Red Hook, NY, USA: Curran Associates Inc., 26548–26560.

Lamour J, Serbin SP, Rogers A, Acebron KT, Ainsworth E, Albert LP, Alonzo M, Anderson J, Atkin OK, Barbier N, et al. 2026. The Global Spectra-Trait Initiative: A database of paired leaf spectroscopy and functional traits associated with leaf photosynthetic capacity. Earth System Science Data 18: 245–265.

Leek JT, Scharpf RB, Bravo HC, Simcha D, Langmead B, Johnson WE, Geman D, Baggerly K, Irizarry RA. 2010. Tackling the widespread and critical impact of batch effects in high-throughput data. Nature Reviews Genetics 11: 733–739.

Lochocki EB, Salesse-Smith CE, McGrath JM. 2025. PhotoGEA: An R Package for Closer Fitting of Photosynthetic Gas Exchange Data With Non-Gaussian Confidence Interval Estimation. Plant, Cell & Environment 48: 5104–5119.

Ma Y, Chen S, Ermon S, Lobell DB. 2024. Transfer learning in environmental remote sensing. Remote Sensing of Environment 301: 113924.

McGreivy N, Hakim A. 2024. Weak baselines and reporting biases lead to overoptimism in machine learning for fluid-related partial differential equations. Nature Machine Intelligence 6: 1256–1269.

Meacham-Hensold K, Montes CM, Wu J, Guan K, Fu P, Ainsworth EA, Pederson T, Moore CE, Brown KL, Raines C, et al. 2019. High-throughput field phenotyping using hyperspectral reflectance and partial least squares regression (PLSR) reveals genetic modifications to photosynthetic capacity. Remote Sensing of Environment 231: 111176.

Meireles JE, Cavender-Bares J, Townsend PA, Ustin S, Gamon JA, Schweiger AK, Schaepman ME, Asner GP, Martin RE, Singh A, et al. 2020. Leaf reflectance spectra capture the evolutionary history of seed plants. New Phytologist 228: 485–493.

Meyer H, Pebesma E. 2022. Machine learning-based global maps of ecological variables and the challenge of assessing them. Nature Communications 13: 2208.

Ng W, Minasny B, Montazerolghaem M, Padarian J, Ferguson R, Bailey S, McBratney AB. 2019. Convolutional neural network for simultaneous prediction of several soil properties using visible/near-infrared, mid-infrared, and their combined spectra. Geoderma 352: 251– 267.

Paoletti ME, Haut JM, Plaza J, Plaza A. 2019. Deep learning classifiers for hyperspectral imaging: A review. ISPRS Journal of Photogrammetry and Remote Sensing 158: 279–317.

Passos D. 2026. Convolutional neural networks in Vis–NIR chemometrics: From contradiction to conditional design. TrAC Trends in Analytical Chemistry 203: 119015.

Ploton P, Mortier F, Réjou-Méchain M, Barbier N, Picard N, Rossi V, Dormann C, Cornu G, Viennois G, Bayol N, et al. 2020. Spatial validation reveals poor predictive performance of large-scale ecological mapping models. Nature Communications 11: 4540.

Rackauckas C, Ma Y, Martensen J, Warner C, Zubov K, Supekar R, Skinner D, Ramadhan A, Edelman A. 2021. Universal Differential Equations for Scientific Machine Learning.

Raue A, Kreutz C, Maiwald T, Bachmann J, Schilling M, Klingmüller U, Timmer J. 2009. Structural and practical identifiability analysis of partially observed dynamical models by exploiting the profile likelihood. Bioinformatics 25: 1923–1929.

Reichstein M, Camps-Valls G, Stevens B, Jung M, Denzler J, Carvalhais N, Prabhat. 2019. Deep learning and process understanding for data-driven Earth system science. Nature 566: 195–204.

Riedlinger A, Kretschmer J, Möller K. 2015. On the practical identifiability of a two-parameter model of pulmonary gas exchange. BioMedical Engineering OnLine 14: 82.

Roberts DR, Bahn V, Ciuti S, Boyce MS, Elith J, Guillera-Arroita G, Hauenstein S, Lahoz-Monfort JJ, Schröder B, Thuiller W, et al. 2017. Cross-validation strategies for data with temporal, spatial, hierarchical, or phylogenetic structure. Ecography 40: 913–929.

Rogers A. 2014. The use and misuse of Vc,max in Earth System Models. Photosynthesis Research 119: 15–29.

Rogers A, Serbin SP, Ely KS, Sloan VL, Wullschleger SD. 2017. Terrestrial biosphere models underestimate photosynthetic capacity and CO2 assimilation in the Arctic. New Phytologist 216: 1090–1103.

Serbin SP, Dillaway DN, Kruger EL, Townsend PA. 2012. Leaf optical properties reflect variation in photosynthetic metabolism and its sensitivity to temperature. Journal of Experimental Botany 63: 489–502.

Sharkey TD. 2019. Is triose phosphate utilization important for understanding photosynthesis? Journal of Experimental Botany 70: 5521–5525.

Shen C, Appling AP, Gentine P, Bandai T, Gupta H, Tartakovsky A, Baity-Jesi M, Fenicia F, Kifer D, Li L, et al. 2023. Differentiable modelling to unify machine learning and physical models for geosciences. Nature Reviews Earth & Environment 4: 552–567.

Smith NG, Keenan TF, Colin Prentice I, Wang H, Wright IJ, Niinemets Ü, Crous KY, Domingues TF, Guerrieri R, Yoko Ishida F, et al. 2019. Global photosynthetic capacity is optimized to the environment. Ecology Letters 22: 506–517.

Song G, Wang Q. 2021. Including Leaf Traits Improves a Deep Neural Network Model for Predicting Photosynthetic Capacity from Reflectance. Remote Sensing 13: 4467.

Stinziano JR, McDermitt DK, Lynch DJ, Saathoff AJ, Morgan PB, Hanson DT. 2019. The rapid A/Ci response: a guide to best practices. New Phytologist 221: 625–627.

Stock A, Gregr EJ, Chan KMA. 2023. Data leakage jeopardizes ecological applications of machine learning. Nature Ecology & Evolution 7: 1743–1745.

Valavi R, Elith J, Lahoz-Monfort JJ, Guillera-Arroita G. 2019. blockCV: An r package for generating spatially or environmentally separated folds for k-fold cross-validation of species distribution models. Methods in Ecology and Evolution 10: 225–232.

Verrelst J, Malenovský Z, Van der Tol C, Camps-Valls G, Gastellu-Etchegorry J-P, Lewis P, North P, Moreno J. 2019. Quantifying Vegetation Biophysical Variables from Imaging Spectroscopy Data: A Review on Retrieval Methods. Surveys in Geophysics 40: 589–629.

Walker AP, Quaife T, van Bodegom PM, De Kauwe MG, Keenan TF, Joiner J, Lomas MR, MacBean N, Xu C, Yang X, et al. 2017. The impact of alternative trait-scaling hypotheses for the maximum photosynthetic carboxylation rate (Vcmax) on global gross primary production. New Phytologist 215: 1370–1386.

Wei Y, Ye Z, Jiang C, Bernacchi CJ, Liang C, Zhao G, Yao N, Fan X, Yu Q, Wu G. 2026. Leaf CO2 response curve fitting method matters: cross-tool assumptions spread photosynthetic parameters and bias leaf–canopy photosynthesis. New Phytologist 251: 2427–2445.

Xu R, Ferguson J, Breil-Aubert M, Kromdijk J, Nikoloski Z. 2026. Generalizability and transferability of machine learning models using hyperspectral reflectance data for maize traits. Scientific Reports 16: 5865.

Yendrek CR, Tomaz T, Montes CM, Cao Y, Morse AM, Brown PJ, McIntyre LM, Leakey ADB, Ainsworth EA. 2017. High-Throughput Phenotyping of Maize Leaf Physiological and Biochemical Traits Using Hyperspectral Reflectance. Plant Physiology 173: 614–626.

Yin X, Amthor JS. 2024. Estimating leaf day respiration from conventional gas exchange measurements. New Phytologist 241: 52–58.

Zhu XX, Tuia D, Mou L, Xia G-S, Zhang L, Xu F, Fraundorfer F. 2017. Deep Learning in Remote Sensing: A Comprehensive Review and List of Resources. IEEE Geoscience and Remote Sensing Magazine 5: 8–36.

